# A carbonic anhydrase pseudogene sensitizes select *Brucella* lineages to low CO_2_ tension

**DOI:** 10.1101/721597

**Authors:** Lydia M. Varesio, Jonathan W. Willett, Aretha Fiebig, Sean Crosson

## Abstract

*Brucella* are intracellular pathogens that cause a disease known as brucellosis. Though the genus is highly monomorphic at the genetic level, species have animal host preferences and some defining physiologic characteristics. Of note is the requirement for increased CO_2_ tension to cultivate particular species, which confounded early efforts to isolate *B. abortus* from diseased cattle. Differences in the capacity of *Brucella* species to assimilate CO_2_ are determined by mutations in the carbonic anhydrase gene, *bcaA*. Ancestral single nucleotide insertions in *bcaA* have resulted in frameshifted pseudogenes in *B. abortus* and *B. ovis* lineages, which underlie their inability to grow under the low CO_2_ tension of a standard atmosphere. Incubation of wild-type *B. ovis* in air selects for mutations that “rescue” a functional *bcaA* reading frame, which enables growth under low CO_2_ and enhances growth rate in high CO_2_. Consistent with this result, we show that heterologous expression of functional *E. coli* carbonic anhydrases enables *B. ovis* growth in air. Growth of wild-type *B. ovis* is acutely sensitive to a reduction in CO_2_ tension, while frame-rescued *B. ovis* mutants are completely insensitive to CO_2_ shifts. Wild-type *B. ovis* initiates a gene expression program upon CO_2_ downshift that resembles the stringent response and results in activation of its *virB* type IV secretion system. Our study provides evidence that loss-of-function insertion mutations in *bcaA* sensitize the response of *B. ovis* and *B. abortus* to reduced CO_2_ tension relative to other *Brucella* lineages. CO_2_-dependent starvation and virulence gene expression programs in these species may influence persistence or transmission in natural hosts.

**Importance:** *Brucella* spp. are highly related, but exhibit differences in animal host preference that must be determined by genome sequence differences. *B. ovis* and the majority of *B. abortus* strains require increased CO_2_ tension to be cultivated *in vitro*, and harbor conserved insertional mutations in the carbonic anhydrase, *bcaA*, which underlie this trait. Mutants that grow in a standard atmosphere, first reported nearly a century ago, are easily selected in the laboratory. These mutants harbor varied indel polymorphisms in *bcaA* that restore its consensus reading frame and rescue its function. Loss of *bcaA* function has evolved independently in the *B. ovis* and *B. abortus* lineages, and results in a dramatically increased sensitivity to CO_2_ limitation.

## Introduction

Bacteria of the genus *Brucella* are the etiologic agents of brucellosis, which is among the most common zoonotic diseases worldwide (1). *Brucella* spp. can infect a range of wild and livestock animals (2), but have a relatively narrow host range and varying zoonotic potential (3, 4) and pathogenicity in humans. *B. melitensis*, for example, is primarily a sheep and goat pathogen and is often considered the most virulent species in humans (5). In contrast, human infections by the cattle pathogen, *B. abortus*, and the swine pathogen, *B. suis* (biovars 1, 3, and 4), are less frequent and often less pathogenic, though clinical differences between these species are difficult to discern (6). Considering that genetic identity across the genus is 94-98% at the coding level (4, 7), differences in *Brucella* spp. host range, virulence, and zoonotic potential are notable. Comparative studies of genome content and the genetic requirements for growth in diverse environments may inform our understanding of differences in the physiology and infection biology of *Brucella* spp.

A historically-significant physiologic difference between species is the requirement for CO_2_ supplementation for axenic growth. In this manuscript, we provide a historical overview of CO_2_ as a required substrate for *Brucella* cultivation, describe a series of forward and reverse genetic and transcriptomic studies focused on understanding CO_2_ metabolism in the ovine pathogen, *Brucella ovis*, and present a genus-level evolutionary analysis of a β-carbonic anhydrase gene (*bcaA*) that provides understanding of why select *Brucella* species require CO_2_ supplementation for axenic growth.

### Brucella *cultivation and CO*_*2*_: *a brief history*

Bang and Stribolt pioneered the study of *B. abortus* growth in axenic culture, which was important in establishing its identity as the etiologic agent of contagious abortion in cattle (8). Of particular note in their original study was the growth pattern of *B. abortus* in solid shake tubes, in which they observed bacteria in two bands below the surface of nutrient agar. Based on a series of experiments using different atmospheres of CO_2_ and O_2_, Bang concluded that oxygen and not carbon dioxide was the gaseous factor that governed growth, though he noted significant variability in growth zones in shake tubes (8). In the years following the discovery of Bang and Stribolt, reliable axenic cultivation of *B. abortus* from clinical specimens remained a major challenge (9).

Huddleson conducted a series of growth experiments in 1921, demonstrating that increasing the partial pressure of CO_2_ (pCO_2_) stimulated growth of *B. abortus* clinical isolates (10). A follow-up investigation provided additional evidence that CO_2_ was required to support metabolism of *B. abortus*, and that the effect of CO_2_ on growth was not a result of pH changes in the medium (11). G.S. Wilson later presented evidence that both oxygen and CO_2_ *per se* are required for *B. abortus* growth, and that bicarbonate cannot substitute for CO_2_ in the growth medium (12).

To assess the metabolic fate of CO_2_, J.B. Wilson and colleagues grew multiple *B. abortus* strains in the presence of ^14^CO_2_ and measured incorporation of carbon into amino acids and nucleotides (13-15). Carbon-14 largely pooled in pyrimidines and in glycine: 90% of the carbon-14 detected in the amino acid pool was in glycine (15). Amino acids, purines, pyrimidines, various organic acids and cell hydrolysates were not able to substitute for CO_2_ in their experiments (16). In the case of *B. abortus*, the incorporation of ^14^CO_2_ into glycine and pyrimidines occurred at similar ratios in all strains tested, even those that did not require increased pCO_2_ for growth (15). From these data, the authors concluded that fixation of CO_2_ is a fundamental property of *B. abortus* strains.

### Natural variation in the requirement for CO_2_ supplementation

When investigating the cause of Malta fever, Bruce noted that *Micrococcus* (*Brucella*) *melitensis* appeared as colonies “on the surface” of nutrient agar in an air incubator (17). This stands in contrast with *Brucella abortus* isolates from cattle that did not grow on agar surfaces (8) without CO_2_ supplementation (10). Indeed, variation in atmospheric requirements for growth of these bacteria clouded the fact that these early investigators were actually studying closely-related species (18). Pioneering work by Alice Evans united the etiologic agents of Malta fever in humans and contagious abortion in cattle into a single group (19) that she classified under the genus *Brucella* (20). By 1930, it was established that certain *Brucella* isolates required CO_2_ supplementation for axenic cultivation, while others could be cultivated without increased pCO_2_ (21). In the case of *B. abortus*, it was well known by the 1920’s that the requirement for increased pCO_2_ for growth was often spontaneously lost after a period of cultivation in the laboratory (11, 22). Buddle reported a similar phenomenon in an early description of *Brucella ovis* isolated from rams in New Zealand (23).

What, then, occurs when a *Brucella* isolate spontaneously loses the requirement for increased pCO_2_ to grow? Early descriptions of “crops” of colonies that grew from clinical specimens on solid media after several days of incubation (11) are reminiscent of spontaneous genetic mutants that arise when bacteria are cultivated under selective conditions. Smith attempted to isolate genetic revertants of lab-adapted strains derived from these crops by exposing the cells to guinea pig tissue or re-infecting cows (11). However, he failed to identify strains that regained a requirement for CO_2_ supplementation. Marr and Wilson later demonstrated that *B. abortus* spontaneously loses its requirement for CO_2_ at a rate of 3 × 10^-10^ per generation (24), supporting a model whereby this phenotype is likely a result of mutation at a single site.

### A model to investigate Brucella responses to CO_2_ limitation

As a model to study CO_2_ metabolism and gene regulation, we have selected *Brucella ovis*, a natural pathogen of sheep, that has a well-documented requirement for increased pCO_2_ to grow axenically (23, 25, 26). We report a forward genetic selection strategy to identify *B. ovis* mutants that grow spontaneously under standard atmospheric conditions, and have defined the genetic changes in these strains by whole genome sequencing. In agreement with recent results from Perez-Etayo and colleagues (27), we found that *B. ovis* mutants selected to grow in a standard atmosphere carry a frameshift mutation at the 3’ end of a β-carbonic anhydrase gene, *bcaA*_*BOV*._ Carbonic anhydrases catalyze the reversible hydration of CO_2_ to bicarbonate, an essential metabolic building block (28), and plasmid-based expression of the frameshifted *bcaA*_*BOV*_ allele or of functional *Escherichia coli* carbonic anhydrases, *can* or *cynT*, are sufficient to enable growth of *B. ovis* in a standard atmosphere.

To better understand the selected *bcaA*_*BOV*_ frameshift mutations, we conducted a comparative and evolutionary analysis of over 700 sequenced *Brucella* genomes, and identified polymorphisms in *B. ovis* and select *B. abortus* strains that result in non-functional, frameshifted *bcaA* pseudogenes. Our data provide evidence that *B. ovis* mutants selected to grow without CO_2_ supplementation have “resurrected” the frameshifted *bcaA*_*BOV*_ pseudogene to encode a functional BcaA protein sequence. The protein sequence of the resurrected BcaA_BOV_ matches BcaA from *B. melitensis* and other *Brucella* species that do not require CO_2_ supplementation. From these data, we conclude that the different atmospheric requirements for *Brucella* spp. cultivation noted by early investigators of *Brucella* physiology are a result of small insertion and deletion mutations in *bcaA*, and that functional revertants of *bcaA* pseudogenes are easily selected under CO_2_ limitation. We further show that *1)* wild-type *B. ovis* is acutely sensitive to CO_2_ limitation as a result of harboring a *bcaA* pseudogene, *2)* that CO_2_ limitation triggers transcription of known starvation stress response and virulence genes including the *virB* type IV secretion system, and *3)* that gain-of-function frameshift mutations in *bcaA* render *B. ovis* insensitive to changes in CO_2_ tension.

## Materials and Methods

### Bacterial strains and growth conditions

The bacterial strains used in this study are listed in **Data Set 4**. *Brucella ovis* was grown on Schaedler agar (Difco Laboratories) or on Tryptic Soy Agar (Difco Laboratories) plates supplemented with 5% defibrinated sheep blood (Quad Five) (SBA or TSBA respectively) at 37°C with 5% CO_2_ when required. Liquid cultures were grown in Brucella Broth (BB) (Difco Laboratories) at 37°C with 5% CO_2_ when required. 50 µg ml^-1^ of kanamycin (kan) was added when required. Expression strains were supplemented with 1 mM isopropy β-D-1-thiogalactopyranoside (IPTG) (GoldBio). All studies on *Brucella ovis* were performed following Biosafety Level 2 (BSL2) protocols.

*Escherichia coli* strains used for cloning were grown on lysogeny broth (LB) (Fisher Bioreagents) at 37°C and on LB plates with 1.5% agar (Fisher Bioreagents). Kan was added to a final concentration of 50 µg ml^-1^. For conjugation, the diaminopimelic acid (DAP) auxotrophic *E. coli* strain WM3064 was grown with a final concentration of 300 µM DAP (Sigma-Aldrich).

### Plasmid and Strain Construction

For the construction of allele replacement strains, external primers overlapping regions approximately 500 bp upstream or downstream flanking target genes of interest were PCR amplified with KOD Xtreme Hot Start Polymerase (Novagen) using either *Brucella ovis, Brucella ovis* mutant or *Brucella abortus* genomic DNA as a template. For deletion strains, internal primers were designed so that the 4-6 amino acids at the N-terminus and the C-terminus of the gene product would remain in frame to minimize polar effects. For strains with point or small mutations, the target gene was amplified using internal primers carrying the mutation of interest. The various fragments obtained were then joined by overlap extension PCR (OE-PCR) to generate null or mutant alleles using KOD. Insertion of fragments into the suicide plasmid pNPTS138 was attained either by restriction enzyme (RE) digestion and T4 ligase ligation or by Gibson assembly (New England Biolabs).

To build gene overexpression strains, genomic DNA from *Escherichia coli* MG1655, *Brucella ovis, Brucella abortus* or derivatives thereof was used as templates. Primers amplifying the gene of interest starting with the start codon and including the stop codon were amplified by PCR with KOD. Fragment insertion into the *lac*-inducible, low-copy pSRK plasmid was attained either by RE digestion and T4 ligation or by Gibson assembly. All insertion sequences into plasmids were confirmed by Sanger sequencing. Sequence-confirmed plasmids were transformed into *E. coli* Top10 (for plasmid maintenance), purified (ThermoFisher Scientific), and transformed into *E. coli* WM3064 (William Metcalf, University of Illinois) for conjugation into *Brucella ovis*. In the case of replicating plasmids, selection on SBA or TSBA kan plates (without DAP) allowed for isolation of single *B. ovis* clones containing the plasmid of interest. In the case of the pNPTS138 plasmid used to conduct gene replacement, cells from mating were first selected on SBA or TSBA kan plates. pNPTS138 also carries the *sacB* gene for counterselection. Strains harboring pNPTS138 integrants were outgrown in BB and spread on TSBA (or SBA) plates containing 5% (w/v) sucrose. This permitted selection of single clones in which a second recombination event that excised the plasmid had occurred. Clones were then screened by PCR with GoTaq® Green polymerase and the PCR products were Sanger sequenced (University of Chicago Comprehensive Cancer Center, DNA Sequencing & Genotyping Facility) to confirm the sequence of the gene deletion or replacement. See **Data Set 4** for primers, strains and plasmids.

### DNA extraction, amplification and quantification

Genomic DNA was extracted following a standard guanidinium thiocyanate protocol. Briefly, strains were struck on SBA (or TSBA) agar plates and colonies were picked and cultivated in BB overnight. 1 ml of culture was spun at 12000 rpm for 20 s and the pellet washed with 0.5 ml of Phosphate-Buffered Saline (PBS). Pellet was resuspended in 0.1 ml of TE buffer pH 8.0 (10mM Tris-HCl pH 8.0 (Fisher Bioreagents); 1 mM EDTA (Fisher Bioreagents)). TE buffer included ribonuclease A (1 µl ml^-1^). 0.5 ml of GES lysis solution (5 M guanidinium thiocyanate (Fisher Bioreagents), 0.5 M EDTA pH 8.0; 0.5% v/v sarkosyl) was added. Following a 15 min incubation at 60 °C, 0.25 ml of cold 7.5 M ammonium acetate (Fisher Bioreagents) were added. After 10 min on ice, 0.5 ml of chloroform (Fisher Bioreagents) were added and samples were vortexed and centrifuged. The aqueous phase was mixed with 0.54 volumes of cold isopropanol (Fisher Bioreagents) and incubated at room temperature for 15 min before centrifugation. The pellet was washed three times in 70% ethanol before resuspending in TE buffer + RNAseA, at which point the concentration was determined spectrophotometrically.

### Forward genetic selection for mutants that grow without added CO_2_

To select for spontaneous mutants that grow without addition of 5% CO_2_ to the atmosphere wild-type *B. ovis* ATCC 25840 or *B. ovis* Δ*bcaA* strains were inoculated into BB at a final optical density (600 nm; OD_600_) of ∼0.3 then placed in a shaker (200 rpm) under standard atmospheric conditions (0.04% CO_2_). Cultures were incubated at 37°C and spectrophotometrically monitored for evidence of growth every 24 hours (**Fig. S1A)**. A total of 29 independent spontaneous mutants were selected that acquired the ability to grow without added CO_2_. These mutant isolates were back diluted to OD_600_=1.5 × 10^-3^, 1.5 × 10^-5^, 1.5 × 10^-7^, 1.5 × 10^-9^ and grown in 0.04% CO_2_ to confirm that the isolates indeed grew without the addition of 5% CO_2_ (result from inoculum of 1.5 ×10^-5^ shown in **Table S1**). Samples were confirmed to *B. ovis* by PCR amplification of a *Brucella*-specific gene (BOV_RS05580).

### Whole Genome DNA Sequencing (WGS)

Genomic DNA (gDNA) from the parent *B. ovis* strain, which requires 5% CO_2_ to grow, and gDNA from 16 independent spontaneous mutants that grow without added CO_2_ (evolved from the parent stock) was purified using the standard guanidinium thiocyanate-based procedure described above. DNA sequencing libraries were prepared from randomly sheared DNA and sequenced using the standard Illumina protocol (Illumina HiSeq 4000; single end 50 bp reads). Reads were mapped to the *Brucella ovis* ATCC 25840 genome (chromosome 1 and chromosome 2 RefSeq accession numbers NC_009505 and NC_009504, respectively) and polymorphisms were identified using breseq (72) (https://github.com/barricklab/breseq). Raw DNA sequencing reads for all strains are available in the NCBI Sequence Read Archive (SRA), through BioProject accession PRJNA540707.

### Growth Curves

Strains were struck on SBA or TSBA plates and allowed to grow for ∼48 hours. Initial inoculum into BB ranged from OD_600_ of 0.015 to 0.03. Antibiotics for plasmid maintenance and IPTG for expression induction were added when needed at the start of the growth experiment. Growth of two to three separate tubes as technical replicates per sample was monitored spectrophotometrically at 600 nm (OD_600_). Each growth curve was measured at least three times independently.

### Sequence alignments

Genome sequences of *Brucella* isolates were downloaded from PATRIC (https://www.patricbrc.org/) (73). The amino acid sequence was aligned using the multiple alignment tool in Geneious 10.0.9 (www.geneious.com): global alignment with free end gaps, Blosum 62 scoring matrix, gap open penalty of 12 and gap extension penalty of 3. To cluster the *Brucella abortus bcaA* alleles, nucleotide sequences were aligned using Geneious multiple alignment, global alignment with free end gaps specifically, cost matrix set at 65% similarity, gap open penalty of 12, gap extension penalty of 3 and 2 refinement iterations.

### RNA sample preparation and RNA sequencing

Samples for RNA sequencing were prepared as follows (see also **Fig. S3A**): six independent cultures each of wild-type *B. ovis* or *B. ovis bcaA1*_*BOV*_ strain were inoculated into BB and allowed to grow to saturation overnight at 37°C in a roller in the presence of 5% CO_2_. Cells were back diluted to an OD_600_ of 0.05 and monitored until they reached OD_600_ of 0.1. Three tubes of *B. ovis bcaA1*_*BOV*_ were transferred to a roller in a standard air incubator (i.e. 0.04% CO_2_), while three tubes of *bcaA1*_*BOV*_ remained in the 5% CO_2_ incubator. Both sets of *bcaA1*_*BOV*_ cultures were harvested after an additional 2.5 hours of growth. Cells from three tubes of wild-type *B. ovis* were harvested at this same time, while the last three tubes of wild-type *B. ovis* were transferred to the standard air incubator and rolled for an additional 2.5 hours. Wild-type *B. ovis* cultures showed no signs of growth over this period in air (0.04% CO_2_). Cells from the remaining three wild-type *B. ovis* tubes incubated in the atmospheric (0.04% CO_2_) incubator were then harvested.

RNA was prepped from all harvested cell samples as follows: each individual culture sample was aliquoted into six 1.5 ml Eppendorf tubes (for a total of 9 ml) and spun for 60 s at 14,500 × g. Pelleted cells were immediately re-suspended in 1 ml of Trizol (Ambion, Life Technologies, ThermoFisher Scientific) and stored at - 80 °C. Samples were then thawed at 65 °C for 10 minutes. Once thawed, 200 µl of cold chloroform was added, cells were vortexed for 15 s and incubated for 5 min at room temperature. Samples were then centrifuged at 4 °C (17000 × g) for 4 min and 500 µl of 100% cold isopropanol was added to the clear supernatant in fresh tube, then frozen for at least 1hr at -80 °C. Samples were centrifuged at 4 °C (17000 × g) for 30min, the supernatant was removed, and 1 ml of 70% ethanol was added to wash the pellet. After an additional 5 min centrifuge (at 4 °C, 17000 × g), residual ethanol was removed and the RNA pellet was resuspended in RNAse free water (50 µl). RNA was treated with DNAse and further purified with RNA purification kit (Qiagen). Concentrations were determined spectrophotometrically, and RNA quality was initially assessed by running 1 µl of each sample on a Tris/Borate/EDTA (TBE) 2% agarose gel.

Ribosomal RNA was depleted from the sample using the Gram-Negative bacteria Ribo-Zero rRNA Removal Kit (Illumina-Epicentre). RNA-seq libraries were prepared with an Illumina TruSeq stranded RNA kit according to manufacturer’s instructions. The libraries were sequenced on an Illumina HiSeq 4000. Sequencing reads were deposited in the NCBI GEO database and are available under accession GSE130678.

### RNA sequencing data analysis

Reads were mapped to the *Brucella ovis* ATCC 25840 genome (chromosome I and chromosome II RefSeq accession numbers NC_009505 and NC_009504, respectively) using CLC Genomics Workbench 11.0 (https://www.qiagenbioinformatics.com); mismatch cost: 2; insertion cost: 3; deletion cost: 3; length fraction; 0.8, similarity fraction: 0.8. Two samples were extreme outliers: wild-type *B. ovis* (5% CO_2_) replicate 3 and *B. ovis bcaA1*_*BOV*_ (0.04% CO_2_) replicate 1, based on principle component analysis of the raw expression values. This was likely due to RNA sample degradation in these samples during library preparation. Accordingly, these samples were not included in our differential expression analysis. Reads Per Kilobase per Million (RPKM), Transcripts Per Million (TPM), and Counts Per Million (CPM) values for single samples as well as paired differential expressions – with p-values, False Discovery Rate (FDR) and Bonferroni corrections – are presented in **Data Set 1**. Significance thresholds were arbitrarily set at FDR p-value < 0.001 and fold change > |2|. KEGG analysis of significantly regulated genes in wild-type *B. ovis* in low (0.04%) versus high (5%) CO_2_ conditions was performed using KEGG pathways (74) (https://www.genome.jp/kegg/). KEGG ontology annotations were assigned to these genes. This annotated gene list was submitted to KEGG Mapper–Search Pathways, querying against *Brucella ovis* ATCC 25840 (bov). Pathways that had 2 or fewer genes were removed. We removed from consideration pathways where the number of upregulated versus downregulated genes was not significantly different. Specifically, pathways were removed if the number of the upregulated genes divided by the total number of genes assigned to that pathway was between 0.4 and 0.6.

### Construction and mapping of *B. ovis* ATCC 25840 Tn-Himar mutant library

*B. ovis* ATCC 25840 was struck on an SBA plate and incubated for 48h before mating with *E. coli* APA752 (WM3064 donor strain carrying pKMW3 mariner transposon vector library) (75). *E. coli* APA752 was incubated overnight prior to mating (1 ml was thawed and inoculated into 25 ml of LB + DAP + kan). 10 ml of donor strain were pelleted and mixed with pellet of *B. ovis* ATCC 25840 scraped off an SBA plate at a 1:10 ratio, and resuspended in a total of 250 µl of BB. 50 µl of the conjugation mixture were spotted on SBA + DAP plates and left to incubate overnight. Spots were then harvested and resuspended in 5 ml of BB. Cells were diluted to OD_600_ of 0.032 and 500 µl were plated on each of twelve 150 mm SBA + kan plates. Colonies grew in approximately three days and were harvested and resuspended in 250 ml of BB + kan at a final OD_600_ of 0.2. Cells were allowed to double 2-3 times before being frozen in 15% glycerol. For mapping, the library was thawed and the cells were washed once in PBS before pelleting for DNA extraction. The Illumina sequencing library to map Tn-Himar insertion sites was built following the protocol of Wetmore *et al*. (75) and as previously described (76, 77). Briefly, adapters were added to the genomic library by PCR (30 min at 95 °C and 15 min at 70 °C) using Mod2_TS_Univ and Mod2_TruSeq primers. DNA was then sheared to obtain fragments approximately 300 to 500 bp in length. Samples were then size selected, end repaired, ligated to a custom adapter, and cleaned before 150 bp single end sequencing by the University of Chicago Functional Genomics Facility. See **Table 3** and **Data Set 5** for list of sequencing statistics and unhit genes, respectively.

### Statistical analysis

In bar graphs, error bars represent standard deviation (unless otherwise indicated) of replicates from at least three independent experiments (two or more technical replicates were used for each individual experiment). Strains that failed to grow are indicated as “no growth”. **** indicates significance of p<0.0001, *** indicates significance of p<0.001, ** is p<0.01; ns = non-significant, calculated using one-way ANOVA followed by Tukey’s test using GraphPad Prism version 8.0.2. For the scatter plot, R^2^ values were calculated by implementing linear regression in GraphPad Prism. For heat map, z-scores were calculated as follows: for each gene, the CPM value was multiplied by the mean of CPM values for that gene across all samples. The resulting value was divided by the standard deviation of the CPM values of that gene across all samples. The heat map was built using Java Tree View version 1.1.6r4 (http://jtreeview.sourceforge.net).

## Results

### A forward genetic selection identifies *B. ovis* mutant strains capable of growth in an unsupplemented atmosphere

*Brucella ovis* requires 5% CO_2_ supplementation to grow axenically (25). We sought to identify genes linked to this growth requirement, and thus developed a forward genetic selection to identify spontaneous *B. ovis* mutants that grow in a standard, unsupplemented atmosphere (0.04% CO_2_). Specifically, we inoculated wild-type *B. ovis* in Brucella broth (BB) and allowed the cultures to shake in an air incubator. After several days, some cultures became visibly turbid, providing evidence for growth (see **Fig. S1A** and **Materials and Methods**). Twenty-nine mutant strains were isolated from five independent selections. We inoculated all mutants into fresh broth and confirmed that they indeed grew without CO_2_ supplementation **(Table S1**).

Genomic DNA from 16 of these spontaneous mutants was purified and sequenced along with DNA from the wild-type *B. ovis* parent. Each mutant carried a single nucleotide deletion within a six-nucleotide interval at the 3’ end of a β-carbonic anhydrase gene (locus tag BOV_RS08635; *bcaA*_*BOV*_) (**Fig. 1A and Table 1**). These deletions in *bcaA*_*BOV*_ frameshifted the coding sequence of the BcaA_BOV_ C-terminus and increased the length of the predicted gene product from 207 to 214 residues (**Fig. 1A-C**). The 16 sequenced mutants clustered into 4 groups based on the site of the frameshift mutation; the corresponding alleles are named *bcaA1*_*BOV*_*-4*_*BOV*_ (**Fig. 1C**). Nucleotide deletions in *bcaA*_*BOV*_ were the only mutations that were common to all mutant strains and missing from the parent (**Table 1**).

**Figure 1.**
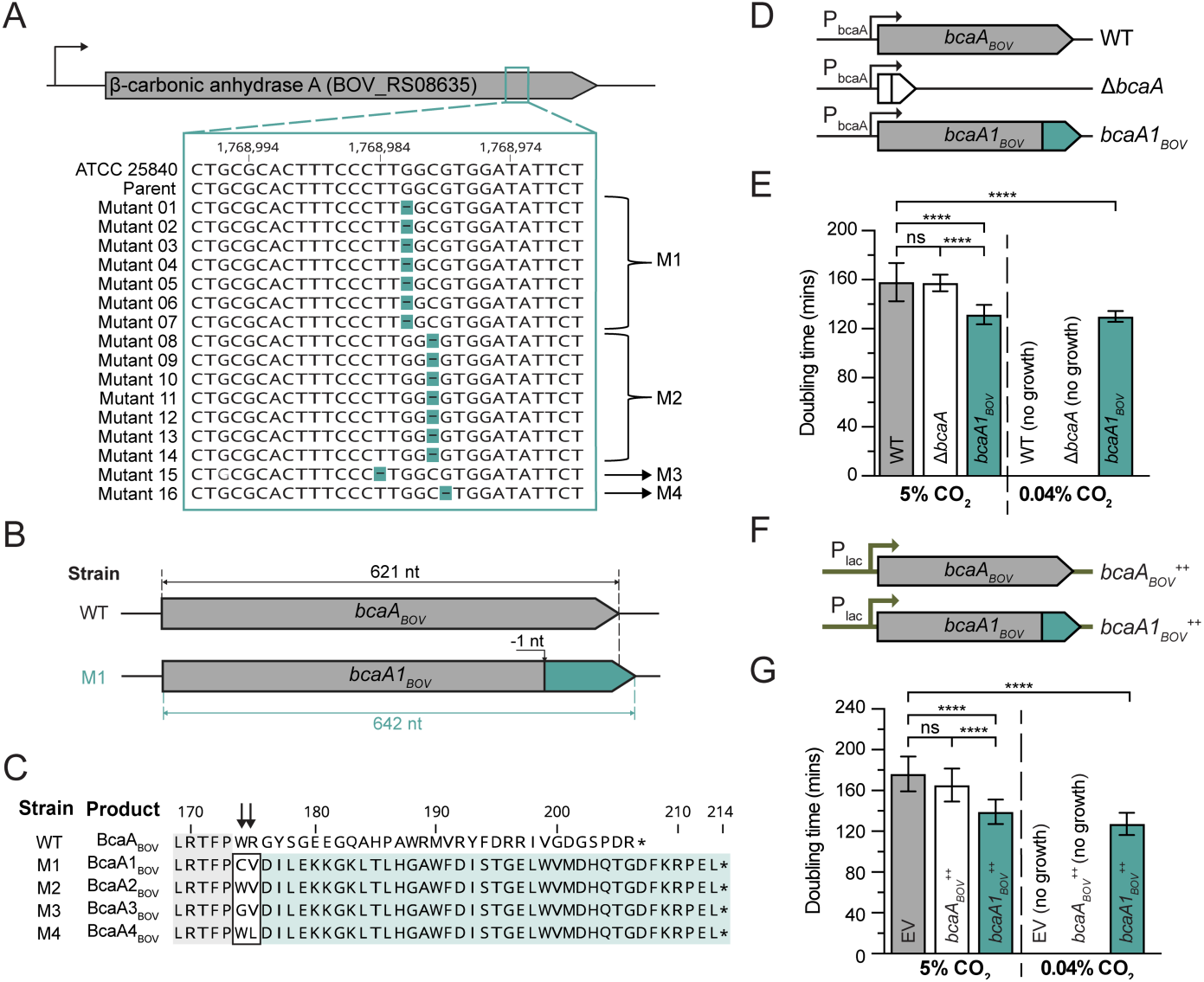
A single nucleotide deletion at the 3’ end of *bcaA*_*BOV*_ enables *B. ovis* growth without CO_2_ supplementation. **A)** Alignment of a segment of the 3’ end of *bcaA*_*BOV*_ (BOV_RS08635) from wild-type *B. ovis* and the 16 selected mutants that can grow without added CO_2_. Gaps show the single nucleotide deletions **(highlighted teal dashes)** in the *bcaA* locus. M1, M2, M3 and M4 are the four classes of single nucleotide deletions that we observed. Annotated nucleotide position is indicated at the top. **B)** Schematic representation of the wild-type *B. ovis* ATCC 25840 allele, *bcaA*_*BOV*_ **(top)**, compared to the frameshifted mutant allele from the M1 cluster of mutants, *bcaA1*_*BOV*_ **(bottom)**. The site of the frameshift is indicated with a vertical arrow. **Teal** shading indicates the portion of the gene that has an altered coding sequence following the frameshift that results from deletion of a single guanosine at position 1,768,986. **C)** Multiple protein sequence alignment of the C-terminal frameshift site of wild-type *B. ovis* ATCC 25840 BcaA_BOV_ and the four selected mutant alleles (M1 to M4). **Teal** shaded area highlights the frameshifted amino acids. **Gray** shaded area highlights sequence that is conserved in all five alleles. **Boxed** amino acids show protein sequence variation at the frameshift site **(black arrows). D)** Schematic of the *bcaA* locus in strains used for experiments in panel **E**; wild-type *B. ovis* ATCC 25480 *bcaA*_*BOV*_ allele **(top)**, in-frame *bcaA*_*BOV*_ deletion allele (Δ*bcaA*) **(middle)**, and a *B. ovis* strain in which the wild-type allele is replaced with the frameshifted *bcaA1*_*BOV*_ allele at the native locus **(bottom)**. P_bcaA_ indicates expression from the native *bcaA* promoter. **E)** Bar graph of doubling times in minutes for the three strains in **panel D** grown with either 5% CO_2_ supplementation or in air without added CO_2_ (0.04%). Error bars represent standard deviation of replicates from three independent experiments (each performed with at least two technical replicates), for a total of 8 measurements per sample. Strains that failed to grow are indicated with “no growth”. **** indicates significance of p<0.0001, calculated using one-way ANOVA followed by Tukey’s post-test. **F)** Schematic of RK2-derived plasmids (pSRK) harboring two different *bcaA* alleles. pSRK carrying a *lac* inducible (P_lac_) wild-type *bcaA*_*BOV*_ **(top)** or frameshifted *bcaA1*_*BOV*_ **(bottom)** were transformed into wild-type *B. ovis* ATCC 25840. Plasmid-bearing strains are referred to as *bcaA*_*BOV*_ ++ and *bcaA1*_*BOV*_ ++, respectively. **G)** Bar graph of doubling times (in minutes) of strains carrying pSRK-*bcaA*_*BOV*_ (*bcaA*_*BOV*_ ++), pSRK-*bcaA1*_*BOV*_ (*bcaA1*_*BOV*_ ++) or the empty vector (EV) as control. Cells were induced with 1mM IPTG, grown in BB with kanamycin to maintain plasmids, and cultivated in a standard air incubator (0.04% CO_2_) or with 5% CO_2_ supplementation. Error bars represent standard deviation of four independent experiments executed in triplicate (total number of measurements per sample = 12). Strains that failed to grow are indicated with “no growth”. **** indicates significance of p<0.0001, calculated using one-way ANOVA followed by Tukey’s post-test.

**Table 1.** Sequence polymorphisms^a^ between 16 independent *B. ovis* mutants that grow without CO_2_ supplementation and the wild-type parent strain^b^.

| Mutant | Chromosome | Position | Mutation | Locus<sup>c</sup> | Gene Annotation | Description |
| --- | --- | --- | --- | --- | --- | --- |
| 01 | NC_009505 | 805,096 | C → T | BOV_RS03980 | NADH-quinone oxidoreductase subunit B | T188T (ACC → ACT) |
|  | NC_009505 | 1,768,982 | -C | BOV_RS08635 | β-carbonic anhydrase - *bcaA<sub>BOV</sub>* | *bcaA<sub>BOV</sub>* → *bcaA1<sub>BOV</sub>* (frameshift) |
| 02<sup>d</sup> | NC_009505 | 1,768,982 | -C | BOV_RS08635 | β-carbonic anhydrase - *bcaA<sub>BOV</sub>* | *bcaA<sub>BOV</sub>* → *bcaA1<sub>BOV</sub>* (frameshift) |
| 03 | NC_009505 | 1,768,980 | -G | BOV_RS08635 | β-carbonic anhydrase - *bcaA<sub>BOV</sub>* | *bcaA<sub>BOV</sub>* → *bcaA2<sub>BOV</sub>* (frameshift) |
| 04 | NC_009505 | 1,768,980 | -G | BOV_RS08635 | β-carbonic anhydrase - *bcaA<sub>BOV</sub>* | *bcaA<sub>BOV</sub>* → *bcaA2<sub>BOV</sub>* (frameshift) |
| 05 | NC_009505 | 1,768,982 | -C | BOV_RS08635 | β-carbonic anhydrase - *bcaA<sub>BOV</sub>* | *bcaA<sub>BOV</sub>* → *bcaA1<sub>BOV</sub>* (frameshift) |
| 06 | NC_009505 | 1,768,982 | -C | BOV_RS08635 | β-carbonic anhydrase - *bcaA<sub>BOV</sub>* | *bcaA<sub>BOV</sub>* → *bcaA1<sub>BOV</sub>* (frameshift) |
| 07 | NC_009505 | 1,768,980 | -G | BOV_RS08635 | β-carbonic anhydrase - *bcaA<sub>BOV</sub>* | *bcaA<sub>BOV</sub>* → *bcaA2<sub>BOV</sub>* (frameshift) |
| 08 | NC_009505 | 1,768,984 | -A | BOV_RS08635 | β-carbonic anhydrase - *bcaA<sub>BOV</sub>* | *bcaA<sub>BOV</sub>* → *bcaA3<sub>BOV</sub>* (frameshift) |
| 09 | NC_009505 | 301,445 | A → G | BOV_RS0175 | IS5/IS1182 family transposase | L224P (CTT → CCT) |
|  | NC_009505 | 301,958 | A → C | BOV_RS01475 | sensor histidine kinase | V53G (GTT → GGT) |
|  | NC_009505 | 420,434 | G → T | BOV_RS02035 ← /<br>← BOV_RS16215 | hypothetical protein / hypothetical protein | intergenic |
|  | NC_009505 | 1,725,552 | C → T | BOV_RS08460 | branched-chain amino acid ABC transporter ATP-binding protein | G349D (GGC → GAC) |
|  | NC_009505 | 1,768,980 | -G | BOV_RS08635 | β-carbonic anhydrase - *bcaA<sub>BOV</sub>* | *bcaA<sub>BOV</sub>* → *bcaA2<sub>BOV</sub>* (frameshift) |
|  | NC_009504 | 117,883 | (T)8 →<br>(T)7 | BOV_RS10890 → /<br>→ BOV_RS12585 | HTH-type quorum sensing-dependent transcriptional regulator VjbR | intergenic |
|  | NC_009504 | 676,505 | G → A | BOV_RS13635 | sulfurtransferase FdhD | pseudogene (415/810 nt) |
|  | NC_009504 | 908,237 | C → T | BOV_RS14715 → /<br>→ BOV_RS14720 | alpha-hydroxy-acid oxidizing protein lldD / porin family protein | intergenic |
| 10 | NC_009505 | 805,096 | C → T | BOV_RS03980 | NADH-quinone oxidoreductase subunit B | T188T (ACC → ACT) |
|  | NC_009505 | 1,768,982 | -C | BOV_RS08635 | β-carbonic anhydrase - *bcaA<sub>BOV</sub>* | *bcaA<sub>BOV</sub>* → *bcaA1<sub>BOV</sub>* (frameshift) |
| 11 | NC_009505 | 1,768,982 | -C | BOV_RS08635 | β-carbonic anhydrase - *bcaA<sub>BOV</sub>* | *bcaA<sub>BOV</sub>* → *bcaA1<sub>BOV</sub>* (frameshift) |
| 12<sup>d</sup> | NC_009505 | 1,712,213 | G → T | BOV_RS08400 | polyprenyl synthetase family protein | A16S (GCC → TCC) |
|  | NC_009505 | 1,768,979 | -C | BOV_RS08635 | β-carbonic anhydrase - *bcaA<sub>BOV</sub>* | *bcaA<sub>BOV</sub>* → *bcaA4<sub>BOV</sub>* (frameshift) |
| 13 | NC_009505 | 1,768,980 | -G | BOV_RS08635 | β-carbonic anhydrase - *bcaA<sub>BOV</sub>* | *bcaA<sub>BOV</sub>* → *bcaA2<sub>BOV</sub>* (frameshift) |
| 14 | NC_009505 | 1,768,980 | -G | BOV_RS08635 | β-carbonic anhydrase - *bcaA<sub>BOV</sub>* | *bcaA<sub>BOV</sub>* → *bcaA2<sub>BOV</sub>* (frameshift) |
| 15 | NC_009505 | 1,768,980 | -G | BOV_RS08635 | β-carbonic anhydrase - *bcaA<sub>BOV</sub>* | *bcaA<sub>BOV</sub>* → *bcaA2<sub>BOV</sub>* (frameshift) |
| 16 | NC_009505 | 1,768,982 | -C | BOV_RS08635 | β-carbonic anhydrase - *bcaA<sub>BOV</sub>* | *bcaA<sub>BOV</sub>* → *bcaA1<sub>BOV</sub>* (frameshift) |

<sup>a</sup>All mutations shown in the Table are observed in 100% of the reads from the indicated strain
<sup>b</sup> When comparing the sequence of the *B. ovis* ATCC 25840 strain deposited in GenBank to our ATCC 25840 parent strain, 4 differences were present. In 100% of the reads, there was an insertion of a single G in BOV\_RS01360 on chromosome I (NC\_009505), at 282,040:1; an insertion of a single C in an intergenic region on chromosome II (NC\_009504), at position 20:1; and an insertion of a single G in an intergenic region in position 459,010:1, also on chromosome II. These differences were also present in all mutant strains. In ~56% of the sequence reads from the parental strain, we observed a C → T transition (resulting in a T14I coding change) in BOV\_RS10895, *vjbR* at position 118,034 of chromosome II. Each derived mutant strain carried one of several different *vjbR* polymorphisms, each fixed at 100% of the sequencing reads.
<sup>c</sup>For intergenic polymorphisms, ← or → indicate direction of flanking genes
<sup>d</sup>This strain lacks the insertion in BOV\_RS01360 observed in our parental strain, and thus is identical to the deposited *B. ovis* ATCC 25840 GenBank sequence strain at this locus.

Similar *bcaA*_*BOV*_ mutations in *B. ovis* strains that grow without CO_2_ supplementation have been recently described by Perez-Etayo *et al*.(27), though they have used the name carbonic anhydrase II (CAII) for the gene locus and product. For the purposes of this manuscript, we have chosen to use the bacterial genetic naming guidelines of Demerec *et al*. (29), and thus refer to this gene as *bcaA*_*BOV*_. We conclude that cultivation of wild-type *B. ovis* in a standard atmosphere selects for a frameshift mutation at the 3’ end of the β-carbonic anhydrase gene, *bcaA*_*BOV*_. This frameshift increases the length of the predicted protein and completely changes the sequence of the last 32-33 residues of BcaA_BOV_. Based on the fact that we successfully selected mutants in standard air after several days of incubation, we presume that wild-type *B. ovis* can replicate (albeit very slowly) in an unsupplemented atmosphere.

### A *bcaA* frameshift enables growth in an unsupplemented atmosphere

To directly test the hypothesis that *bcaA*_*BOV*_ frameshift mutations enable growth in an unsupplemented atmosphere, we built a strain in which wild-type *bcaA*_*BOV*_ was replaced with the frameshifted *bcaA1*_*BOV*_ allele. We also built a strain in which *bcaA*_*BOV*_ was deleted in-frame (Δ*bcaA*) (**Fig. 1D)**. We measured growth of these strains and wild-type *B. ovis* in 5% CO_2_, and in standard atmospheric CO_2_ levels (approx. 0.04%). All three strains grew in the presence of 5% CO_2_, indicating that *bcaA*_*BOV*_ is not required for growth in a CO_2_-supplemented atmosphere. When these strains were incubated with 0.04% CO_2_, only the *bcaA1*_*BOV*_ allele replacement strain showed measurable growth. Notably, the *bcaA1*_*BOV*_ strain grew significantly faster (i.e. had a shorter doubling time) than wild-type and Δ*bcaA* strains in the presence of 5% CO_2_ (**Fig. 1E and Fig. S1B**). From these data, we conclude that frameshift mutations at the 3’ end of *bcaA1*_*BOV*_ enable robust growth of *B. ovis* in an unsupplemented atmosphere and result in faster growth than wild-type in the presence of 5% CO_2_.

### *bcaA* frameshift mutations are the sole genetic mechanism by which *B. ovis* growth is spontaneously rescued under low CO_2_ tension

To test whether mutations in genes other than *bcaA*_*BOV*_ enable *B. ovis* growth in an unsupplemented atmosphere, we repeated the forward genetic selection described above starting with the Δ*bcaA* strain. We incubated Δ*bcaA* in BB at 37°C for over two weeks in an air incubator and never observed turbid cultures. This experiment was repeated 4 times, with approximately 30 inoculated tubes each time, and growth was never observed. This stands in contrast with our initial selection experiment, in which multiple tubes inoculated with wild-type *B. ovis* became turbid over this timescale. As an alternative forward genetic approach, we generated a pool of *B. ovis* Tn-Himar mutants with insertions at over 5 × 104 unique sites (**Table 2)**. This experiment was aimed at identifying insertional mutations that enable growth in a standard atmosphere. We incubated aliquots of this pool in BB for over two days in an air incubator and did not observe growth. Thus, none of the transposon insertions in the pool enabled *B. ovis* growth in the absence of CO_2_ on this short timescale. From these data, we conclude that single nucleotide deletions at the 3’ end of *bcaA*_*BOV*_, which result in a frameshifted gene product, are the sole genetic mechanism by which *B. ovis* spontaneously acquires the ability to grow without added CO_2_ on this experimental timescale.

**Table 2.**
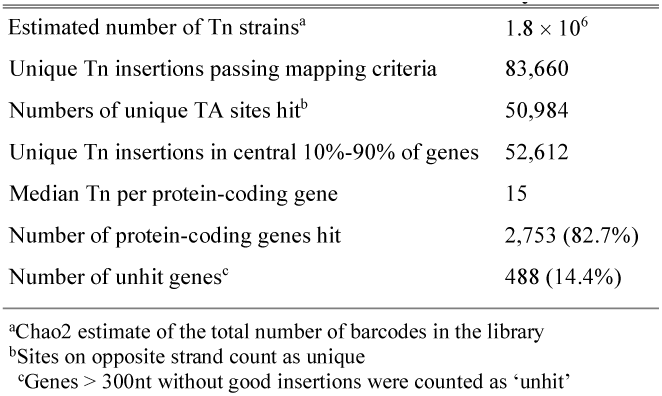
*B. ovis* ATCC 25840 Tn-Himar library.

### The frameshifted alleles, *bcaA1*_*BOV*_*-4*_*BOV*_, are dominant over the wild-type *bcaA*_*BOV*_ allele

We next transformed wild-type *B. ovis* with low-copy replicating plasmids, from which we expressed either the wild-type (*bcaA*_*BOV*_^++^) or frameshifted (*bcaA1*_*BOV*_^++^) alleles from an IPTG-inducible (*lac*) promoter (P_lac_) (**Fig. 1F**). A strain carrying an empty vector (EV) was used as a control. The EV and the *bcaA*_*BOV*_^++^ strains (with 1 mM IPTG) grew comparably in 5% CO_2_ and did not grow when shifted to a standard atmosphere (0.04% CO_2_) (**Fig. 1G and Fig. S1C**). Thus, overexpressing wild-type *bcaA*_*BOV*_ from a plasmid is not sufficient to enable growth in a standard atmosphere. The *bcaA1*_*BOV*_^++^ strain grew without added CO_2_, demonstrating that the frameshifted allele is dominant and sufficient to enable *B. ovis* growth in 0.04% CO_2_. Moreover, the strain expressing *bcaA1*_*BOV*_^++^ grew significantly faster in 5% CO_2_ than the *bcaA*_*BOV*_^++^ and EV strains, providing additional evidence that *bcaA1*_*BOV*_ enhances growth rate in broth culture (**Fig. 1G and Fig. S1C**).

To confirm that all four classes of frameshifted *bcaA* alleles identified in our forward genetic selection (*bcaA1*_*BOV*_*-4*_*BOV*_) were sufficient to enable growth in a standard atmosphere, we expressed these alleles from a replicating plasmid in a standard air incubator (0.04% CO_2_) and compared growth to the *B. ovis* EV strain. As expected, all four alleles enabled growth in 0.04% CO_2_ (**Fig. S1D**). Thus, different single nucleotide deletions at the 3’ end of *bcaA*_*BOV*_ (**Fig. 1C**) result in the same phenotype: growth of *B. ovis* in a standard atmosphere. Given that the four alleles phenocopy each other, we continued our study with the *bcaA1*_*BOV*_ allele. This was the most common isolate from our forward genetic selection and is the most conserved *bcaA* allele in the entire *Brucella* clade (see below).

### Expression of *E. coli* β-carbonic anhydrases is sufficient to enable *B. ovis* growth in a standard atmosphere

*bcaA*_*BOV*_ is predicted to encode a β-class carbonic anhydrase. These enzymes catalyze the reversible hydration of CO_2_ to bicarbonate, an important anabolic substrate. Carbonic anhydrases have been extensively studied (28, 30) in diverse species (31-33), including *Brucella suis* 1330 (34-36). The thirty-three C-terminal residues of wild-type *B. ovis* BcaA_BOV_ differ from *B. suis* 1330 BcaA. However, the sequence of the frameshifted mutant *B. ovis* BcaA1_BOV_ protein is identical to *B. suis* BcaA_BSU1330_ (locus tag: BR_RS08400; also known as _Bs1330_CAII (27) or bsCAII (34)) with the exception of one residue (**Fig. S2A**). The *B. ovis bcaA1*_*BOV*_ mutation thus results in a frameshifted gene product with a C-terminus that matches *B. suis* BcaA_BSU1330_, as well as BcaA orthologs in other *Brucella* species, which we discuss below.

*B. suis* 1330 BcaA_BSU1330_ is reported to be an active β-carbonic anhydrase (34, 35). Thus, we reasoned that the selected frameshift mutations at the 3’ end of *B. ovis bcaA*_*BOV*_ restored the protein to an active form that can hydrate CO_2_ at the low partial CO_2_ pressure of a standard atmosphere, thereby enabling growth. Following this hypothesis, we tested whether heterologous expression of known active *Escherichia coli* MG1655 β-class carbonic anhydrases *can* (locus tag: b0126) or *cynT* (locus tag: b0339) (37) was sufficient to support growth of the wild-type and Δ*bcaA B. ovis* strains in a standard atmosphere **(Figure 2A)**. Induction of *can* or *cynT* expression from a low-copy plasmid (RK2-P_lac_) with 1mM IPTG had no effect on growth rate in 5% CO_2_ (**Fig. 2B**), but enabled growth of wild-type and Δ*bcaA B. ovis* in an unsupplemented atmosphere (0.04% CO_2_) (**Fig. 2C**). We conclude that expression of *E. coli can* or *cynT* (**Fig. S2B**) is sufficient to enable growth of WT and Δ*bcaA B. ovis* strains under the low CO_2_ tension of a standard atmosphere and, therefore, that carbonic anhydrase activity in and of itself is sufficient to allow wild-type or Δ*bcaA B. ovis* to grow in 0.04% CO_2_

**Figure 2.**
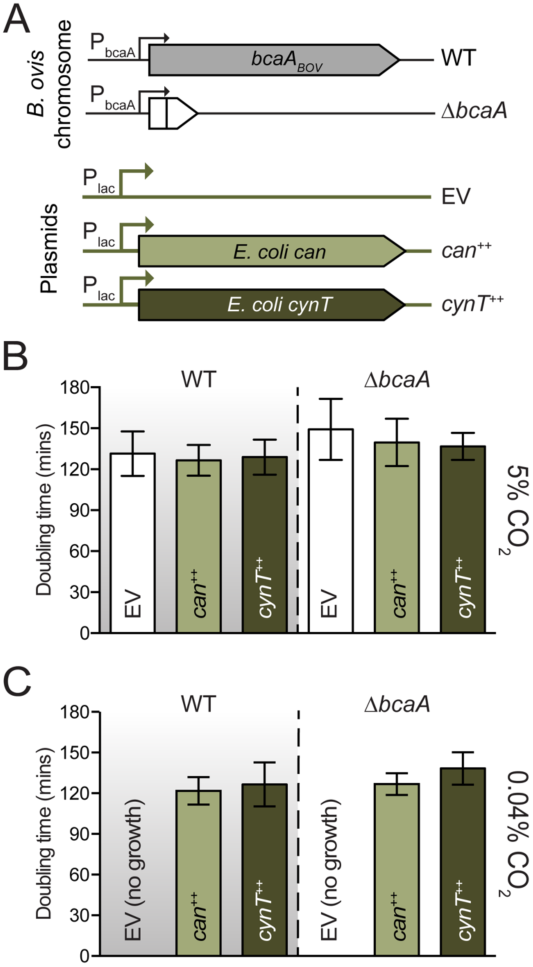
Heterologous expression of two different *Escherichia coli* b-carbonic anhydrases enables growth of wild-type *B. ovis* ATCC 25840 without CO_2_ supplementation. **A)** Schematic representation of RK2-derived plasmids (pSRK) transformed into either wild-type *B. ovis* ATCC 25840 (WT) or the in-frame *bcaA* deletion strain (Δ*bcaA*). pSRK plasmids carried either *E. coli* b-carbonic anhydrase *can* (*can*^++^) or *cynT* (*cynT*^++^) under inducible control of P_lac_. Empty vector (EV) plasmid was used as a control. **B and C)** Doubling time (in minutes) of strains outlined in **A** grown in 5% CO_2_ **(B)** or 0.04% CO_2_ **(C)** after induction with 1mM IPTG. Cells were grown in BB with kanamycin to maintain plasmids. The parent genotype (wild-type or Δ*bcaA*) is indicated at the top of each bar graph. The plasmid genotypes are indicated in each bar. Experiment was performed in triplicate two to four independent times (with two to three technical replicates for a total of 5 to 14 measurements per sample). Error bars represent standard deviation of at least two independent experiments. Strains that failed to grow are indicated with “no growth”.

### A comparative analysis of *bcaA* sequences and the CO_2_ growth requirement in the genus *Brucella*

As outlined in the introduction, *B. ovis* and the majority of *B. abortus* strains require CO_2_ supplementation for axenic cultivation (26). To assess the extent to which variation in *bcaA* is linked to this physiologic trait, we compared the sequences of 773 *bcaA* orthologs from the genus *Brucella* (see **Data Set 1**). *B. ovis* and the majority of *B. abortus* encode BcaA variants that significantly deviate from the consensus sequence for the genus (**Fig. 3A**).

**Figure 3.**
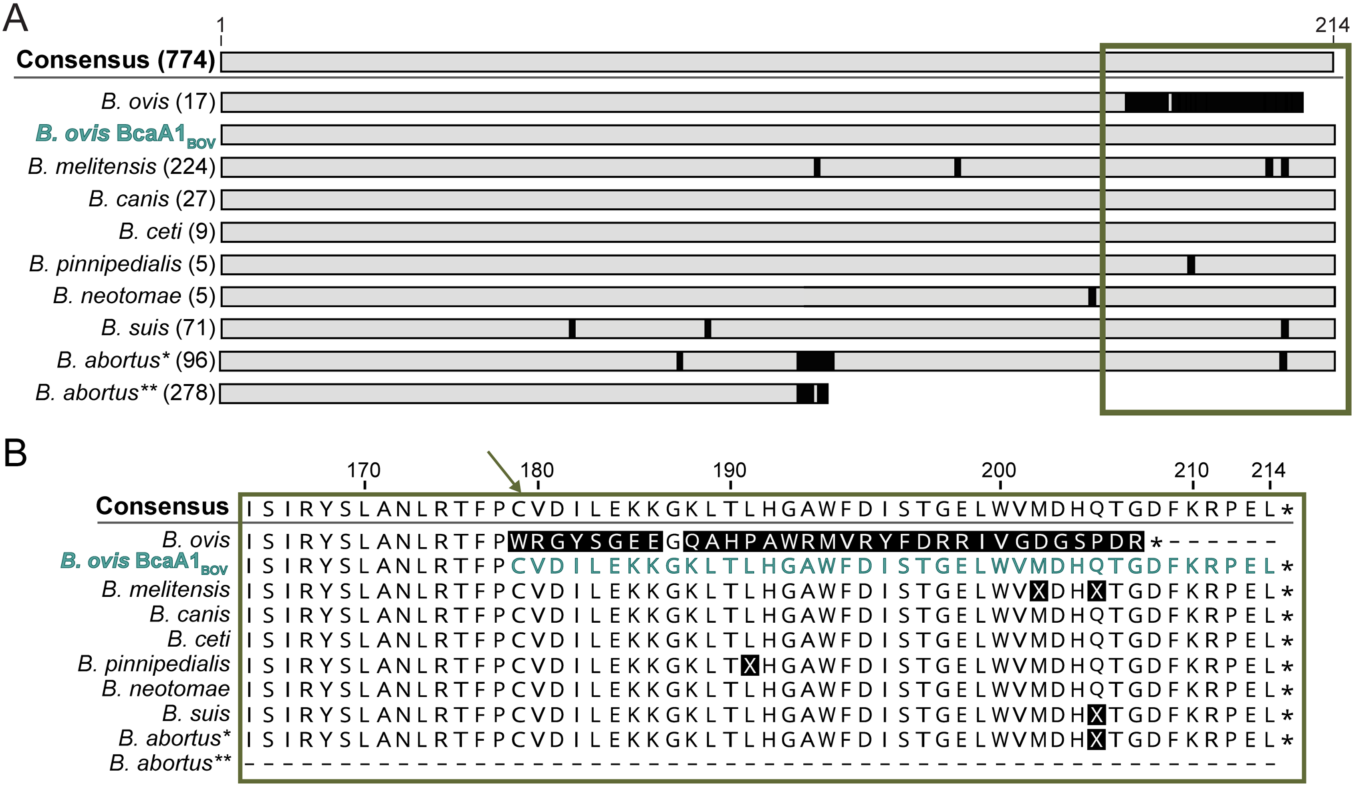
Sequence polymorphisms in BcaA orthologs across the genus *Brucella*. **A)** Schematic of the BcaA protein sequence polymorphisms based on alignment of 774 BcaA orthologs from *Brucella* spp. available in the PATRIC database and found in **Data Set 1**, including *B. ovis* BcaA1_BOV_ (**teal**; M1 cluster). Genus level consensus of all sequences is depicted on top. Species-specific consensus sequences were generated for species with at least 5 representatives (sequences not included in **Fig**.**3** are grouped in ‘Other, **Data set 1**). The species-specific consensuses were then aligned as shown. Amino acid differences between the species-specific and genus-level BcaA consensus sequences are indicated in **black**. *Brucella abortus* sequences were split into two groups, *B. abortus*\* and *B. abortus*\*\*, based on the presence or absence of a frameshift (and premature stop codon) resulting from a single cytosine insertion 339 nucleotides after the annotated start codon, respectively (see also **Fig. 4** and **main text)**. The number of strains compiled in each species-specific consensus sequence is shown in parentheses. The frameshifted BcaA1_BOV_ allele that enables growth of *B. ovis* without added CO_2_ is included in the alignment (**teal**). **B**) Enlargement of green boxed area in (**A**). The C-terminal end of *B. ovis* BcaA1_BOV_ (M1 cluster) is highlighted in **teal**; the site of the frameshift is indicated (**green angled arrow**).

All 17 *B. ovis bcaA*_*BOV*_ sequences in our dataset harbor an additional guanosine 522 nucleotides after the start codon (G522) relative to other *Brucella* species **(Fig. 1)**, which results in a corresponding frameshifted BcaA_BOV_ C-terminus (**Figs. 1 & 3**). The *B. ovis* sequences we analyzed were from strains isolated from sheep in Europe, South America, Australia, and New Zealand over the course of several decades (**Data Set 1**). Thus, this single nucleotide insertion in *bcaA*_*BOV*_ is apparently a defining genetic characteristic of *B. ovis*. As highlighted in **Figs. 1A-C and 3B**, single nucleotide deletion mutations that enable *B. ovis* growth in a standard atmosphere (*bcaA1*_*BOV*_*-4*_*BOV*_) abolish this frameshift and restore the BcaA C-terminus to the consensus of the *Brucella* genus.

We further analyzed *bcaA* from 374 *B. abortus* strains and observed two major sequence classes: *1)* the majority of sequences (278) encode a *bcaA* pseudogene (*B. abortus*\*\* in **Fig. 3**), and *2)* the remainder encode full-length proteins (*B. abortus*\* in **Fig. 3**) that completely or almost completely match consensus (see below). The *bcaA* pseudogene (*bcaA*_*BAB*_) sequences all share the same C339 insertion that results in a frameshift and premature stop codon (**Fig. 4A, cluster 1**). The *bcaA* sequences from the remaining 96 *B. abortus* strains, encoding a full-length BcaA protein, grouped into seven clusters. The largest of these clusters contains sequences lacking the C339 insertion (52 out of 374 strains), resulting in a consensus BcaA gene product **(Figs. 3 and 4A, cluster 2)**. In the other *B. abortus bcaA* clusters the consensus reading frame was restored by either a single nucleotide deletion or two-nucleotide insertions within 12 base pairs (downstream) of the C339 frameshift site **(Fig. 4A)**. We posit that the *bcaA*_*BAB*_ pseudogene is the ancestral allele in *B. abortus*, and that insertion/deletion (i.e. indel) mutations near nucleotide 339 have been selected in different *B. abortus* strains/biovars to resurrect functional BcaA variants that enable growth under low partial CO_2_ pressure.

**Figure 4.**
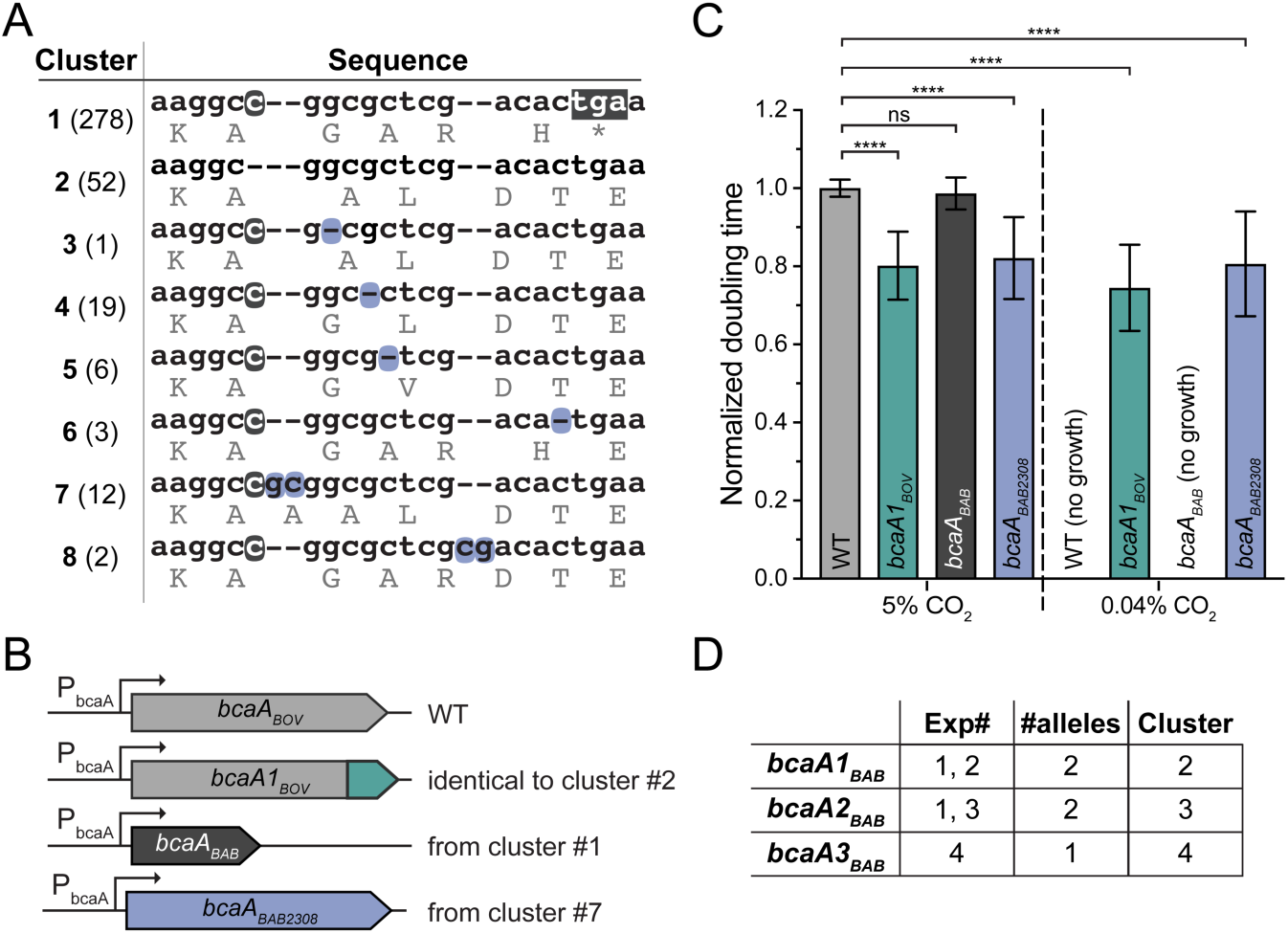
Analysis of sequence and function of *B. abortus bcaA* alleles in *B. ovis* ATCC 25840. **A)** Nucleotide alignment of *B. abortus bcaA* from 374 sequenced isolates starting at position +339 (codon 112). *B. abortus bcaA* sequences cluster into 8 groups based on nucleotide variation. Cluster number is indicated on the left and the number of sequences in each cluster is shown in parenthesis. Translated sequence for each codon is shown in light gray. Cytosine insertion that frameshifts the genus-level consensus (cluster 2) is highlighted **gray. Gray box** in cluster 1 (i.e. allele *bcaA*_*BAB*_) highlights the premature stop codon arising from C insertion. Second-site, singlenucleotide deletions or double nucleotide insertions (highlighted in **blue**) that restore reading frame to consensus are highlighted for clusters 3 through 8. Cluster 7 represents the *bcaA* allele present in *Brucella abortus* strain 2308 (*bcaA*_*BAB2308*_). Clusters 2 and 3 have the same amino acid sequence, but different nt sequence. **B)** Schematic of *bcaA* locus in *B. ovis* ATCC 25840 strains that were functionally assayed in **C**. From **top**: wild-type *bcaA*_*BOV*_ allele, the gain-of-function *bcaA1*_*BOV*_ allele, the *bcaA*_*BAB*_ pseudogene (cluster 1), or the *bcaA*_*BAB2308*_ allele (cluster 7), each expressed from the native promoter (P_bcaA_). **C)** Normalized *B. ovis* doubling times comparing WT (*bcaA*_*BOV*_ allele) to strains harboring *bcaA1*_*BOV*_, *bcaA*_*BAB*_, or *bcaA*_*BAB2308*_ in either high (5%) or low (0.04%) CO_2_. Error bars represent standard deviation of replicates from three to eight independent experiments for a total of 11 to 23 measurements per strain. **** indicates significance of p<0.0001, ns = non-significant, calculated using one-way ANOVA followed by Tukey’s post-test. Strains that failed to grow are indicated as ‘no growth’. **D)** Tabulation of gain-of-function *bcaA*_*BAB*_ mutants that arose in 4 independent selection experiments. Table shows experiment number (Exp#), number of times a specific allele (*bcaA1*_*BAB*_-*3*_*BAB*_) was isolated across the four independent experiments (#alleles), and cluster assignment for that allele (Cluster). Cluster designation as per panel **A**.

### Functional *B. abortus bcaA* alleles are rapidly selected from a *bcaA*_*BAB*_ pseudogene under CO_2_ limitation

We next assayed the function of two *B. abortus bcaA* alleles (*bcaA*_*BAB*_ and *bcaA*_*BAB2308*_) in *B. ovis* (**Fig. 4B**). Specifically, we replaced wild-type *B. ovis bcaA*_*BOV*_ with either the *bcaA*_*BAB*_ pseudogene or the near-consensus *bcaA*_*BAB2308*_ allele (cluster 7, **Figure 4A**), and measured growth in a standard unsupplemented atmosphere (0.04% CO_2_). Replacing *bcaA*_*BOV*_ with *bcaA*_*BAB2308*_ enabled growth of *B. ovis* in 0.04% CO_2_ (**Fig. 4C**), consistent with Perez-Etayo *et al*. (27). Replacing *bcaA*_*BOV*_ with the frameshifted *bcaA*_*BAB*_ pseudogene did not enable *B. ovis* growth in a standard atmosphere (**Fig. 4C**). We conclude that *bcaA*_*BAB*_ is inactive *in vivo* and that the majority of *Brucella abortus* isolates cannot grow in an unsupplemented atmosphere, which is consistent with historical observations outlined in our introduction (8, 10, 11, 22). Conversely, approximately one-quarter of sequenced *B. abortus* strains harbor a copy of *bcaA* in which the frameshift mutation at C339 has apparently been remediated **(Fig. 4A)** to restore expression of a full-length protein. Given these results, we hypothesized that functional variants of *B. abortus bcaA*_*BAB*_ could be experimentally selected in *Brucella* cells cultivated under standard atmospheric conditions, i.e. low pCO_2_.

To test this hypothesis, we used the *B. ovis* strain in which we replaced *bcaA*_*BOV*_ with the non-functional *B. abortus bcaA*_*BAB*_ pseudogene (carrying the C339 insertion) and selected for mutants that could grow in an unsupplemented atmosphere. Four independent experiments yielded spontaneous mutants that grew to high density without CO_2_ supplementation within several days. We sequenced the *bcaA* locus of 5 independent mutants from these selections and identified three types of mutations (*bcaA1*_*BAB*_ *-3*_*BAB*_, **Fig. 4D**) that remediated the C339 frameshift and restored the full-length, consensus BcaA coding frame. All *bcaA* variants that we selected experimentally are present in sequenced isolates of *B. abortus*. We cataloged information of the 374 sequenced *B. abortus* isolates, including assigned biovars (when available) from PATRIC (**Table 3)**. *B. abortus* biovars 5 and 6 have been described as CO_2_ independent (26) and, as predicted, all sequenced isolates of these biovars possess consensus (i.e. full-length) *bcaA* alleles. Biovars 1, 3 and 4 are described as primarily CO_2_ dependent, which is consistent with the fact that between 70 and 95% of sequenced isolates from these biovars carry the *bcaA*_*BAB*_ pseudogene. *B. abortus* biovar 2 is categorized as CO_2_ dependent (26), yet 5 of 18 sequenced isolates possess a consensus functional *bcaA* allele; biovar 7 is classified as CO_2_ independent, though one of five sequenced isolates carries the *bcaA*_*BAB*_ pseudogene. We conclude that the identity of *bcaA* alleles in the different *B. abortus* biovars is largely consistent with the CO_2_ dependence/independence cataloged by Alton *et al*. (26).

**Table 3.**
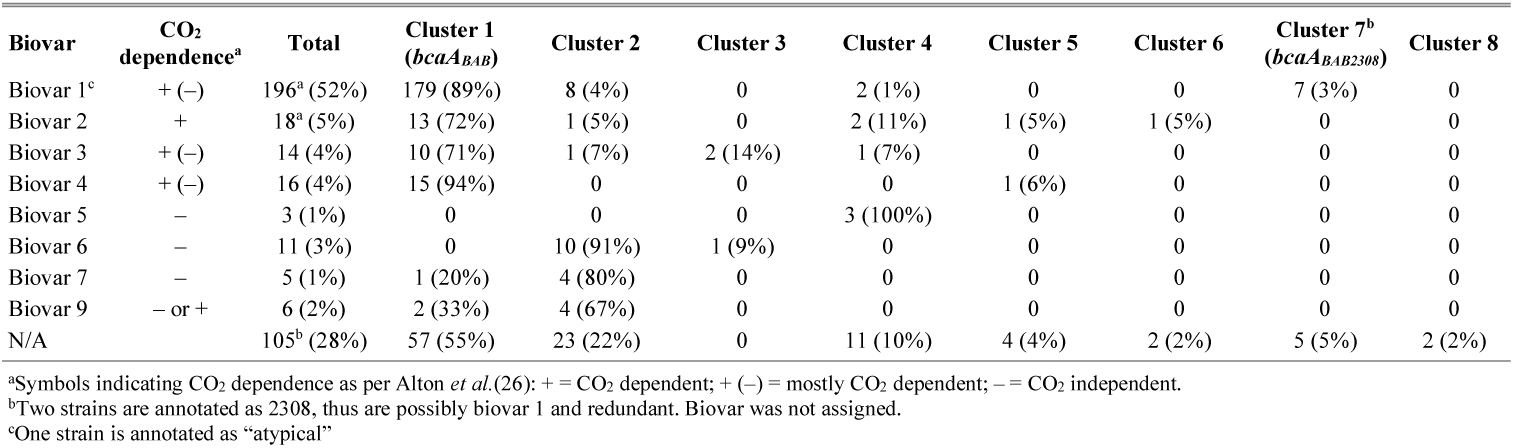
*B. abortus* biovar association with *bcaA* allele clusters.

Our analysis shows that the most common allele of *bcaA* in *B. abortus* is the frameshifted *bcaA*_*BAB*_ pseudogene. Thus, like *B. ovis*, many wild *B. abortus* populations apparently harbor a non-functional, frameshifted *bcaA* pseudogene that renders the bacterium unable to grow without CO_2_ supplementation. Also, like *B. ovis*, functional (consensus) *bcaA* alleles can be easily selected from *bcaA*_*BAB*_ by incubating the bacterium in a standard atmosphere for several days.

### CO_2_ limitation triggers a large-scale transcriptional response in wild-type *B. ovis*, while transcription in the *bcaA1*_*BOV*_ strain is insensitive to CO_2_ shifts

Our group and others (27) have shown that *bcaA* is a specific genetic determinant of growth under low pCO_2_. We have further shown that *B. ovis* and most *B. abortus* strains harbor non-functional *bcaA* pseudogenes containing distinct single-base insertion mutations that result in a frameshifted gene product. These insertional mutations underlie the long-noted CO_2_ requirement for axenic cultivation of these species (see **Introduction**). Given these results, we sought to better understand the physiological consequences of CO_2_ limitation in *B. ovis*, a species that requires CO_2_ for axenic cultivation. Specifically, we defined transcriptional changes in *B. ovis* ATCC 25840 at the genome scale in response a downshift in pCO_2_.

To measure the transcriptional response to CO_2_ change, we cultivated *B. ovis* wild-type and *bcaA1*_*BOV*_ strains in a 5% CO_2_ atmosphere. A subset of these cultures was then shifted to a standard atmosphere (0.04% CO_2_). After a controlled incubation period in this condition (see **Fig. S3A** and **Materials and Methods**) we harvested cells, extracted RNA from all samples, and prepared libraries for sequencing. Shifting wild-type *B. ovis* from 5% to 0.04% CO_2_ slowed growth (**Fig. 5A**), and induced large-scale changes in gene expression: 382 genes were significantly upregulated and 442 genes were significantly downregulated upon a downshift in pCO_2_ (log_2_ fold change > |1|; false-discovery rate (FDR) p-value ≤ 0.001) **(Data Set 2 and Fig. 5B)**. In contrast, shifting a *B. ovis* strain harboring the *bcaA1*_*BOV*_ allele from 5% to 0.04% CO_2_ had no effect on growth **(Fig. 5C**) or gene expression using the same significance threshold **(Data Set 2 and Fig. S3B)**. In fact, a comparison of wild-type *B. ovis* cultivated at 5% CO_2_ to *bcaA1*_*BOV*_ cultivated in 0.04% CO_2_ revealed no differentially expressed genes. As expected from these results, the overall transcriptional profile of wild-type *B. ovis* cultivated in 5% CO_2_ is highly correlated with the *B. ovis bcaA1*_*BOV*_ strain cultivated under either 5% CO_2_ or 0.04% CO_2_ (R^2^= 0.995, 5% CO_2_ and R^2^= 0.993, 0.04% CO_2_) (**Fig. 5D and Fig. S3C)**. We conclude that wild-type *B. ovis* is acutely sensitive to downshifts in levels of atmospheric CO_2_, and that presence of *bcaA1*_*BOV*_ – the consensus allele in the *Brucella* clade – renders *B. ovis* insensitive to CO_2_ downshifts. Notably, our data show that transcription of *bcaA*_*BOV*_ itself is not induced by CO_2_ limitation (**Fig. 5E**).

**Figure 5.**
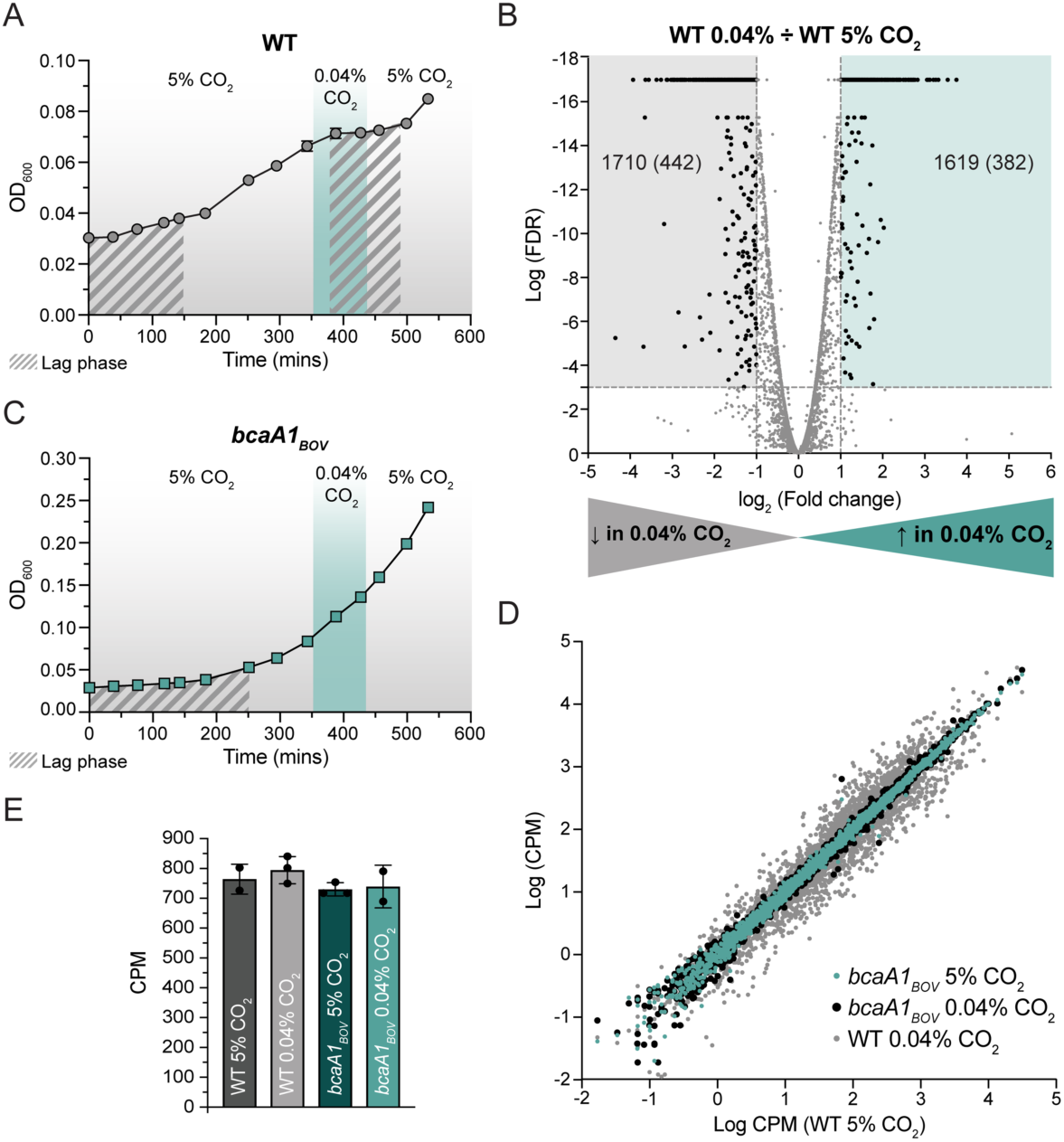
Gene expression changes in wild-type *B. ovis* and *B. ovis bcaA1*_*BOV*_ upon CO_2_ downshift from 5% to 0.04%. **A)** Growth of wild-type *Brucella ovis* ATCC 25840 inoculated in BB and grown in 5% CO_2_. In log phase, cells were shifted to 0.04% CO_2_ (**teal shading**) for approximately 90 minutes then shifted back to 5% CO_2_. Lag phase is highlighted with **gray striped shading**. Experiment was performed three independent times and a representative experiment is shown. **B)** Volcano plot showing log_2_ fold change in gene expression versus log_10_ of the false discovery rate (FDR)-corrected p-value after downshift of wild-type cells from 5% to 0.04% CO_2_. Each dot represents a gene. Genes with FDR p-values < 0.001 and with log_2_ fold changes > |1| were considered statistically significant and are indicated as **black dots**. Due to floating point value constraints in calculation of p-values, the smallest possible FDR p-value we were able to calculate is 1 × 10^-15.^ All lower FDR p-values were arbitrarily assigned a value of 1 × 10^-17^. Genes that were considered upregulated upon CO_2_ downshift are indicated on the right (**teal shading**) and genes that are downregulated upon downshift are indicated on the left (**gray shading**); statistically significant regulated genes in each set are in parenthesis. **C)** Same growth experiment in **(A)** but with the *B. ovis bcaA1*_*BOV*_ strain, which was selected to grow without added CO_2_. **D)** Absolute expression of all measurable transcripts, in log_10_ counts per million (CPM), from wild-type *B. ovis* ATCC 25840 in high (5%) CO_2_ compared to *B. ovis bcaA1*_*BOV*_ in 5% CO_2_ (**teal**), *B. ovis bcaA1*_*BOV*_ in low (0.04%) CO_2_ **(black)** or wild-type in low CO_2_ (**gray**). Each dot represents the mean CPM for a gene (see **Data Set 1**). R^2^ values were calculated (see **main text** and **Fig. S3C**) implementing linear regression using GraphPad Prism 8. CPM were calculated using CLC Genomics Workbench 11.0. **E)** Steady-state *bcaA* transcript levels by strain and condition as indicated. Error bars represent standard deviation calculated from two to three independent samples subjected to RNA-seq measurements.

### CO_2_ limitation induces a stringent-like starvation response and activates transcription of the *virB* type IV secretion gene cluster in wild-type *B. ovis*

We next assigned each of the genes up- or down-regulated upon pCO_2_ downshift in wild-type *B. ovis* to Kyoto Encyclopedia of Genes and Genomes (KEGG) categories. Of the 824 genes that had a log_2_ fold change > |1| and an FDR p-value < 0.001, 447 could be classified into one or more KEGG categories (**Data Set 3)**. Only categories where 3 or more genes were assigned were considered in our pathway analysis (**Fig. 6)**. Genes involved in translation, including those encoding ribosomal proteins, ribosomal RNA, transfer RNA, and aminoacyl-tRNA synthetases, are largely downregulated upon shift from 5% to 0.04% CO_2_ (**Data Set 2, 3 and Fig. 6**). In addition, transcript levels for elongation factors P, Tu and G are highly reduced upon CO_2_ downshift. This transcriptional profile, which reflects reduced protein synthesis, is consistent with nutrient limitation and activation of the stringent response (38-41). In addition, multiple respiratory, energy metabolism, transport, and catabolic genes are downregulated upon CO_2_ downshift **(Data Set 2, 3 and Fig. 6)**. Consistent with the fact that a starvation, or stringent-like response, is induced by CO_2_ downshift, we observe a lag phase in WT *B. ovis* growth upon re-introduction to a 5% CO_2_ environment **(Figure 5A)**.

**Figure 6.**
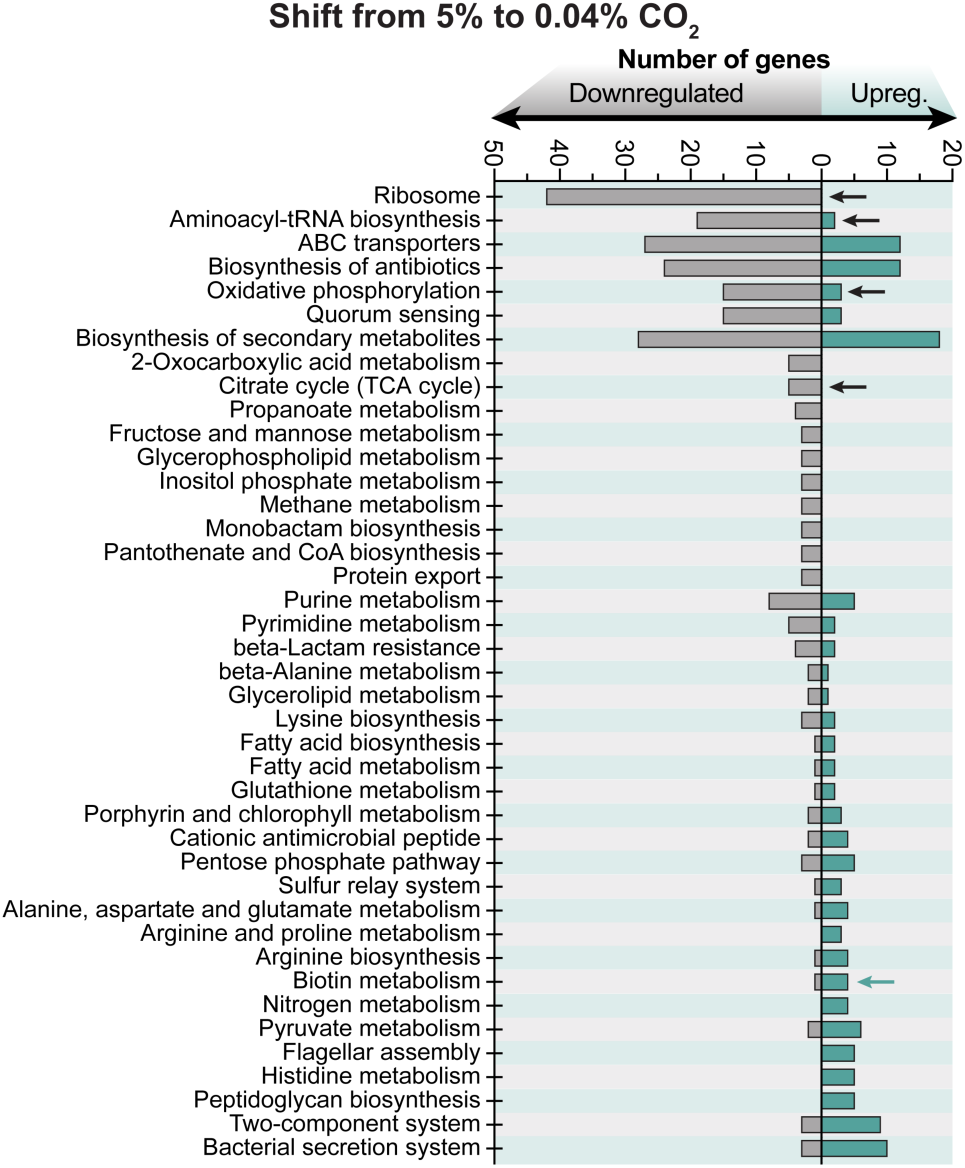
KEGG pathway assignment of genes regulated in wild-type *B. ovis* in response to CO_2_ downshift. KEGG pathways are shown vertically on the left, with bars indicating the number of CO_2_-regulated genes assigned to each pathway. Genes significantly upregulated in response to CO_2_ downshift are on the right (**teal** bars), and genes significantly downregulated in response to CO_2_ downshift are on the left (**gray** bars). Significance thresholds follow those outlined in **Figure 5B** legend. Pathways with less than 3 genes, or where the number of upregulated versus downregulated genes was not significantly different (see **Materials and Methods**), were removed from graph. Select KEGG categories associated with the stringent response are marked by **black arrows. Teal arrow** marks the “Biotin metabolism” pathway.

Expression of genes implicated in response to stress, nutrient limitation, or stationary phase, including polyphosphate kinases (*ppk1, ppk2*) (42-44), small DUF1127-family proteins (*BOV_RS04430, BOV_RS13510, BOV_RS13790, BOV_RS13950*) (45), alternative sigma factors *rpoH* (46) and *ecfG/rpoE1* (47) is activated upon CO_2_ downshift. Furthermore, Entner-Doudoroff/pentose phosphate pathway genes (*zwf, pgl, edd)* are transcriptionally activated, as are genes implicated in *Brucella* spp. virulence and infection including the *virB* type IV secretion gene cluster, the urease cluster, and several annotated flagellar genes **(Figs. 6 & 7 and Data Set 2)**. Although *B. ovis* is urease negative and possesses a degraded urease gene cluster (25), transcription of the *ure* genes (and pseudogenes) is activated upon CO_2_ downshift. Among the most highly activated gene sets in low CO_2_ are biotin biosynthesis genes **(Figs. 6 & 7 and Data Set 2, 3)**. Biotin is a cofactor involved in carboxylation reactions that use CO_2_ as a substrate. Concordant with the result that cells remodel transcription to enhance carboxylation reactions, we also observed that transcription of pyruvate carboxylase and several biotin-dependent acetyl-CoA carboxylases and carboxyl transferases is significantly enhanced **(Fig. 7 and Data Set 2)**.

**Figure 7.**
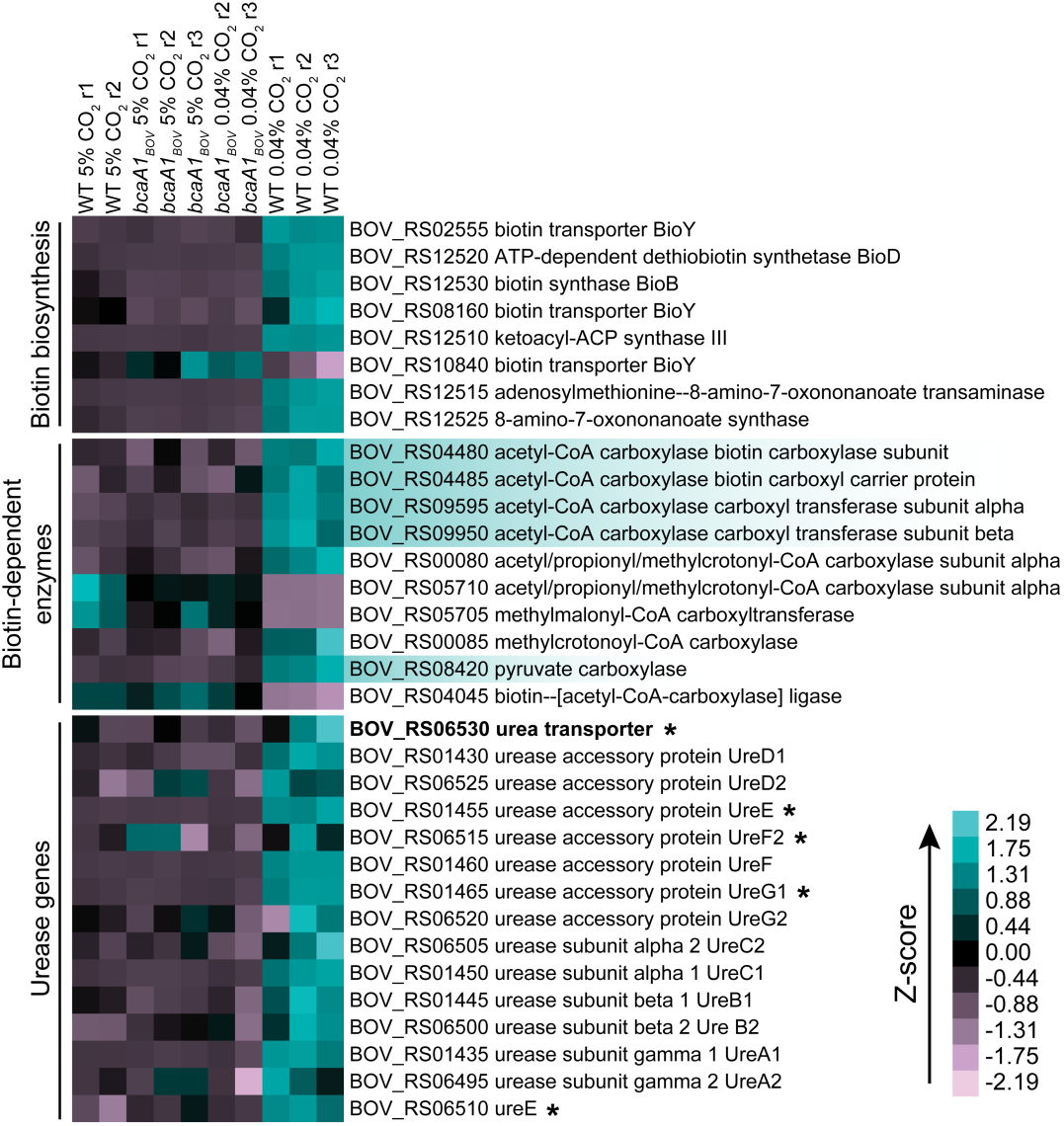
Heat map representation of a subset of genes regulated by CO_2_ downshift in wild-type *B. ovis*. *B. ovis* gene accession numbers and RefSeq annotations are shown to the right of the heatmap. Genes were grouped into three categories: biotin biosynthesis, biotin-dependent enzymes and urease genes. Each column represents an individual RNA sample from either *B. ovis* ATCC 25840 (WT) or *B. ovis bcaA1*_*BOV*_ grown in 5% or downshifted to 0.04% CO_2_, measured by RNA-seq. Strain replicates are indicated as r1, r2, and r3. The four genes that encompass the acetyl-CoA carboxylase complex and pyruvate carboxylase are highlighted in **teal**. *ure* pseudogenes are marked with an asterisk; the urea transporter, which is a pseudogene in *B. ovis*, is in **bold**. Zscores express the number of standard deviations from the mean for each gene (see **Materials and Methods**).

### Pseudogene sequences, including *bcaA*_*BOV*_, are highly conserved across all *B. ovis* isolates

All 17 sequenced *B. ovis* isolates (**Data Set 1**) carry the same single nucleotide insertion in *bcaA*_*BOV*_, yielding a frameshifted gene that apparently encodes a non-functional carbonic anhydrase protein. From these data, we have concluded that *bcaA*_*BOV*_ is a pseudogene. Since pseudogenes are predicted to be under neutral selection (48-50), we might expect that *bcaA*_*BOV*_ should accumulate mutations at a higher frequency than functional genes. Actually, there is evidence that pseudogenes are rapidly deleted from bacterial genomes due to either toxic side effects of expression, or the energetic burden of maintaining the gene (51). However, we observe no other mutations within *bcaA*_*BOV*_ across all sequenced *B. ovis* strains, which have been isolated from regions across the globe over the past 60 years. This suggests either that the nucleotide insertion mutation at the 3’ end of *bcaA*_*BOV*_ occurred very recently in the evolutionary history of the *B. ovis* lineage, or that selective pressures maintain this pseudogene, raising the possibility that this gene sequence may have an additional function. It is known that *bcaA*_*BOV*_ does not contribute to *B. ovis* infection in a mouse model of disease (27), but we have shown that this gene is transcribed **(GEO accession GSE130678 and Fig. 5E)** and thus could contribute other functions. To assess if the conservation of *bcaA*_*BOV*_ allele in *B. ovis* differs from other pseudogenes, we defined the frequency of mutations and sequence divergence across a larger group of both annotated pseudogenes and functional genes, both within in *B. ovis* and compared to *B. abortus* strain 2308. Our aim was to use this gene set to empirically define pseudogene divergence in *B. ovis*, and to use this set as a point of comparison for *bcaA*_*BOV*_.

We quantified polymorphisms in three distinct sets of genes; *1)* urease genes, *2)* the VirB type IV secretion system (T4SS), and *3) B. ovis* pseudogenes described by Tsolis *et al*.(25) (**Table S2**). *B. ovis* is urease negative (23, 52) and the *ure* gene clusters show features of degradation (25) with intact genes and pseudogenes that are presumably both under neutral selection as the pathway is non-functional. *B. abortus*, on the other hand, is urease positive and has an intact *ure* gene cluster (53). The T4SS provided us with a set of genes that are functional and intact in both species (54-56). The additional *B. ovis* pseudogenes included in our analysis are multiple erythritol catabolic genes and cytochrome oxidase genes, which have functional, full-length orthologs in *B. abortus* (**Table S2**). When assessing polymorphisms, each gap (insertion or deletion) was considered as a single event, regardless of its length.

We observed strikingly low divergence within *B. ovis* isolates. In other words, the sequences of all *B. ovis* pseudogenes in our analyzed set are highly conserved across a geographically diverse group of strains isolated between 1959 and 2011. Only 4 pseudogenes — *ureT* (BOV_RS06530), *norB* (BOV_RS11505), *ccoO* (BOV_RS01915) and *pckA* (BOV_RS09880) — exhibit polymorphisms between *B. ovis* strains, and each is polymorphic at only one site **(Table S2)**. Similarly, sequence differences are rare when comparing the *ure* genes or the T4SS genes within the 17 *B. ovis* strains. In short, the level of polymorphism we observe comparing the sequenced *B. ovis* isolates is not significantly different between a functional group of genes (*virB*) and a non-functional gene cluster containing pseudogenes (*ure*).

We then sought to compare these genes to *B. abortus* ATCC 2308. Overall, annotated pseudogenes have the highest average number polymorphisms per base pair between *B. ovis* strains and *B. abortus*. As expected, genes in the functional T4SS (*virB*) locus have the lowest average number of differences per base pair and a bias towards synonymous mutations (**Fig. 8**). *B. ovis* does not produce functional urease and the *ure* genes, accordingly, have a higher number of mutations compared to *B. abortus*. We do observe that larger or out-of-frame deletions are more prevalent in the *ure* genes than the *virB* cluster (**Table S2**) consistent with a model in which the *ure* genes are under relaxed selective pressure. Though pseudogenes are, on average, more highly diverged between *B. ovis* and *B. abortus* (**Fig. 8**), our analysis shows that many other pseudogene sequences besides *bcaA*_*BOV*_ are also entirely conserved in across sequenced *B. ovis* isolates (**Table S2**). Thus, we cannot conclude that the conservation of *bcaA*_*BOV*_ across *B. ovis* isolates is due to purifying selection as a result of BcaA_BOV_ performing some other function in the cell. Rather, a more likely explanation for our results is that the loss of function mutation resulting in a *B. ovis bcaA*_*BOV*_ pseudogene occurred very recently in the evolutionary history of an ancestral *B. ovis* strain that now commonly infects sheep across the globe.

**Figure 8.**
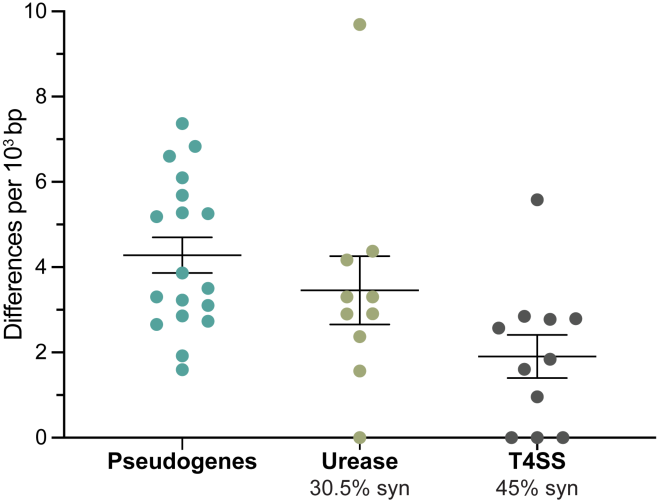
Nucleotide differences between sets of functional genes and pseudogenes in *B. ovis* ATCC 25840 and their *B. abortus* ATCC 2308 orthologs normalized by gene length. Each dot represents a gene. Genes are cataloged in **Table S2**. Bars represent the mean and standard error of the mean for each group. Percentage of synonymous mutations in urease and T4SS genes are indicated

## Discussion

### Brucella cultivation and the resurrection of a carbonic anhydrase pseudogene

During the course of transmission and infection, *Brucella* spp. must endure numerous shifts in the chemical state of the environment (3, 57), including changes in steady-state levels of CO_2_. In the aqueous environment of the cell, CO_2_ can spontaneously hydrate to form bicarbonate, an important substrate for microbial growth (58). The importance of bicarbonate to life is evidenced by the fact that at least 7 different families of carbonic anhydrases (28, 32, 59) have evolved across all kingdoms of life. These enzymes function to enhance the rate of CO_2_ hydration to produce bicarbonate. *Brucella* spp. encode at least three carbonic anhydrases, but our data and recently published data (27) provide evidence that a β-class carbonic anhydrase encoded by *bcaA* is specifically required for axenic growth under the low partial CO_2_ pressure (pCO_2_) of a standard atmosphere in *B. ovis* and *B. abortus*.

Almost all *Brucella* spp. can be cultivated without adding CO_2_, but the majority of *B. abortus* strains and all wild *B. ovis* strains have a strict requirement for CO_2_ supplementation for axenic cultivation. This requirement confounded early efforts to cultivate *B. abortus* from cattle (8, 10), and the knowledge that *B. abortus* strains require added CO_2_ enabled early efforts to cultivate *B. ovis* from sheep tissue (23, 60). Our analysis of over 700 sequenced *Brucella* isolates of multiple species provides evidence that single nucleotide insertions in *bcaA* result in frameshifts that lead to non-functional BcaA protein in *B. abortus* and *B. ovis*. Both of these species carry distinct insertion mutations in *bcaA* (**Figs. 1 & 4 and Data Set 1**) that underlie the strict CO_2_ requirement for growth outside of an animal host. Functional, reverted alleles of these frameshifted *B. abortus* and *B. ovis bcaA* pseudogenes are easily selected by incubating cells in broth under standard atmospheric conditions for several days. This is consistent with historical reports of loss of the CO_2_ requirement after repeated passages of *B. abortus* and *B. ovis* in air (11, 23), and is conceptually in line with a recent report in *E. coli* showing functional resurrection of a pseudogene involved in iron acquisition when cells are cultivated under selective conditions (61).

The fact that *B. ovis* and *B. abortus* lineages have independent insertion/frameshift mutations in *bcaA* that are apparently fixed in populations of each species suggests there may be an adaptive advantage to loss of *bcaA* function. In the case of *B. ovis*, our data show that mutants harboring the functional *bcaA1*_*BOV*_ allele grow significantly faster than wild-type at high partial CO_2_ pressure (**Fig. 1E and 1G**), which is what *Brucella* spp. encounter in mammalian host tissues. The spleen colonization phenotype of *B. ovis* strains carrying a functional *bcaA1*_*BOV*_ allele does not differ from wild-type *B. ovis* in a mouse model of infection (27), but it is known that a slow growth rate can be advantageous for bacterial pathogens to maintain long-term infections in certain tissues (62, 63). Thus, it is possible that loss of *bcaA* function enhances *Brucella* fitness or persistence in tissues of natural (i.e. non-rodent) animal hosts by slowing growth. We note that *B. ovis* is almost entirely restricted to male reproductive tissue and is reported to be transmitted only through direct or venereal contact, via a transiently infected ewe (23, 64, 65). The rate of oxygen uptake and the partial pressure of CO_2_ are higher in testes than in other tissues (66, 67), and it is therefore conceivable that *B. ovis* has evolved a lifestyle in which carbonic anhydrases are no longer required. *B. abortus* has a tropism for bovine reproductive tissue in males and females, and it may be also the case that loss of *bcaA* function influences its rate of growth and its persistence in these tissues, though this hypothesis remains to be tested.

There is no known *Brucella* reservoir outside of animals, but it is well established that presence of bacteria on aborted tissue is one of the main routes of transmission for *B. abortus* (68). A functional BcaA would seem to be advantageous for CO_2_ assimilation when *B. abortus* is residing on such tissue, though the extraordinarily high titer of respiring brucellae on an aborted fetus (69) may lead to a high enough local pCO_2_ that BcaA is unnecessary for growth in this context. Alternatively, *Brucella* growth may be arrested during the window in which it is outside its preferred host tissue. Growth impairment upon CO_2_ limitation may, in fact, serve as an important signal for the bacterium during the transmission cycle. We do not currently understand why loss of *bcaA* function is restricted to the *B. ovis* and *B. abortus* lineages, or if there is a particular feature(s) relating to CO_2_ metabolism or transmission/infection biology that sets these species apart from other members of the genus. It is possible that all *Brucella* species, as intracellular animal pathogens, will eventually lose *bcaA* function.

### CO_2_ limitation elicits a starvation and virulence gene expression response

As outlined in the introduction, carbon-14 from supplied ^14^CO_2_ was recovered primarily in pyrimidines and glycine in *B. abortus* (15, 70). Our efforts to bypass the *B. ovis* CO_2_ requirement by supplementing media with amino acids, nucleotides, and bicarbonate were not successful, just as earlier efforts to bypass the *B. abortus* CO_2_ requirement by adding bicarbonate to the medium were unsuccessful (12). Thus, it seems likely that *Brucella* spp. lack a functional bicarbonate transporter. Our transcriptomic data provide clear evidence that shifting *B. ovis* to a standard atmosphere induces a starvation response that has features of the stringent response (38, 41, 71). This response is evidenced by the downregulation of transcription and translation related genes as well as genes involved in respiration and central metabolism (**Fig. 6**). Among the most significantly activated set of genes upon CO_2_ downshift is the *virB* type IV secretion system, suggesting that cells initiate a gene expression program that could influence virulence when they encounter a CO_2_ limited environment. Growth and transcription in a *B. ovis* strain with a functionally-restored *bcaA* gene (i.e. *bcaA1*_*BOV*_) was completely insensitive to a shift in pCO_2_. We propose that loss-of-function frameshift mutations in *bcaA* yield strains that can more readily sense changes in pCO_2_ in the host. Beyond slowing the growth of *B. ovis* under the high partial CO_2_ pressure of the host environment, it is possible that loss of *bcaA* function also endows cells with an ability to more acutely detect changes in environmental CO_2_ levels, for example when a *Brucella* cell is ejected from its animal host. Whether this change in the CO_2_ detection threshold is advantageous in natural infection and transmission contexts remains to be determined.

## Supporting information

Data Set 1

Data Set 2

Data Set 3

Data Set 4

Data Set 5

## Acknowledgements

We would like to thank Julien Herrou for invaluable experimental help and fruitful scientific discussions. We thank David Hershey for aid in Tn-Himar library construction and helpful scientific input. We also thank the Crosson lab members for the scientific support received. This work was performed with support from the NIH Genetics and Regulation Training Grant (L.M.V.), the NIH R01AI107159 (S.C.), and the Gallo Global Health Fellowship Program (L.M.V.).

## Notes

#### Summary of Updates

This manuscript has been revised based on feedback during the peer review process.

