## Supplemental Figures and Tables for "A carbonic anhydrase pseudogene sensitizes select *Brucella* lineages to low CO_2_ tension"

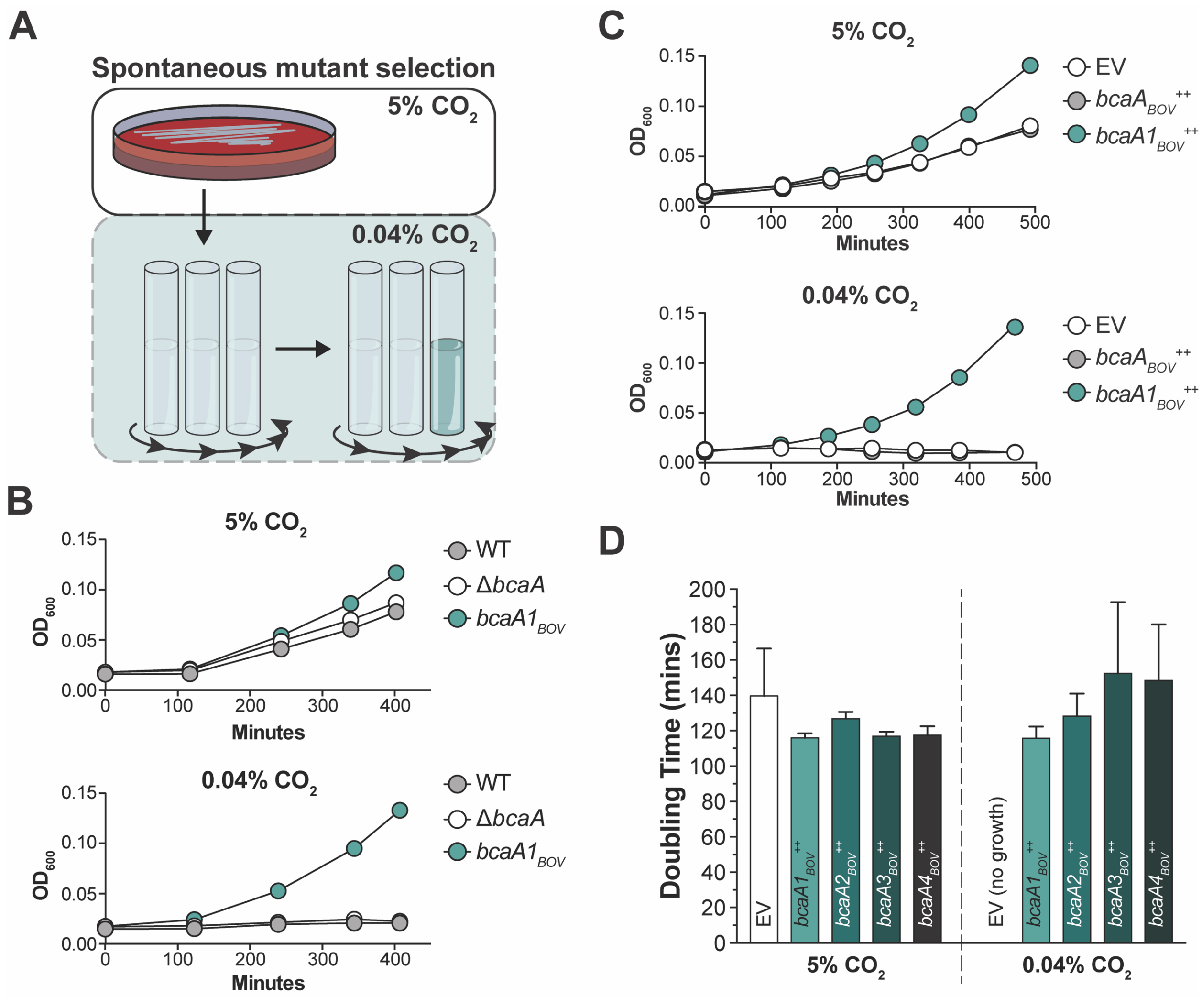


**Supplemental Figure 1. Growth of *B. ovis* harboring *bcaA1$_{BOV}$-4$_{BOV}$* alleles in an unsupplemented atmosphere**
**A)** Cartoon illustrating the forward genetic selection for spontaneous mutants of *Brucella ovis* ATCC 25840 that grow without added $CO_2$. Cells were grown on plates at 37 °C with 5% $CO_2$ supplementation for 48 hrs then inoculated into BB and left in a shaking incubator at 37°C in standard atmospheric conditions (i.e. 0.04% $CO_2$). Growth was monitored by cell culture density measurements ($OD_{600}$) and individual colonies were isolated from tubes where growth was evident. **B)** Growth curve from three independent experiments of *Brucella ovis* ATCC 25840 (WT), Δ*bcaA* and *B. ovis bcaA1$_{BOV}$* strains. Cells were grown either in 5% $CO_2$ (**top**) or 0.04% $CO_2$ (**bottom**). **C)** Growth curve from three independent experiments with *bcaA$_{BOV}$ (bcaA$_{BOV}$$^{++}$*) or *bcaA1$_{BOV}$* (*bcaA1$_{BOV}$ $^{++}$*) overexpressing strains. Cells were grown either in 5% $CO_2$ (**top**) or 0.04% $CO_2$ (**bottom**) after inducing expression from pSRK with 1mM IPTG. Strain carrying the empty vector (EV) was used as a control. **D)** Doubling time of *Brucella ovis* ATCC 25840 strains either carrying the pSRK empty vector plasmid (EV) or one of the four selected ("restored") alleles *bcaA1$_{BOV}$*, *bcaA2$_{BOV}$*, *bcaA3$_{BOV}$* and *bcaA4$_{BOV}$* (*bcaA1$_{BOV}$*$^{++}$, *bcaA2$_{BOV}$*$^{++}$, *bcaA3$_{BOV}$*$^{++}$ and *bcaA4$_{BOV}$*$^{++}$, respectively). Strains that did not grow are indicated (no growth). Experiment was performed on separate days with technical replicates. Bar graphs indicate the standard deviation for 12 measurements per strain.

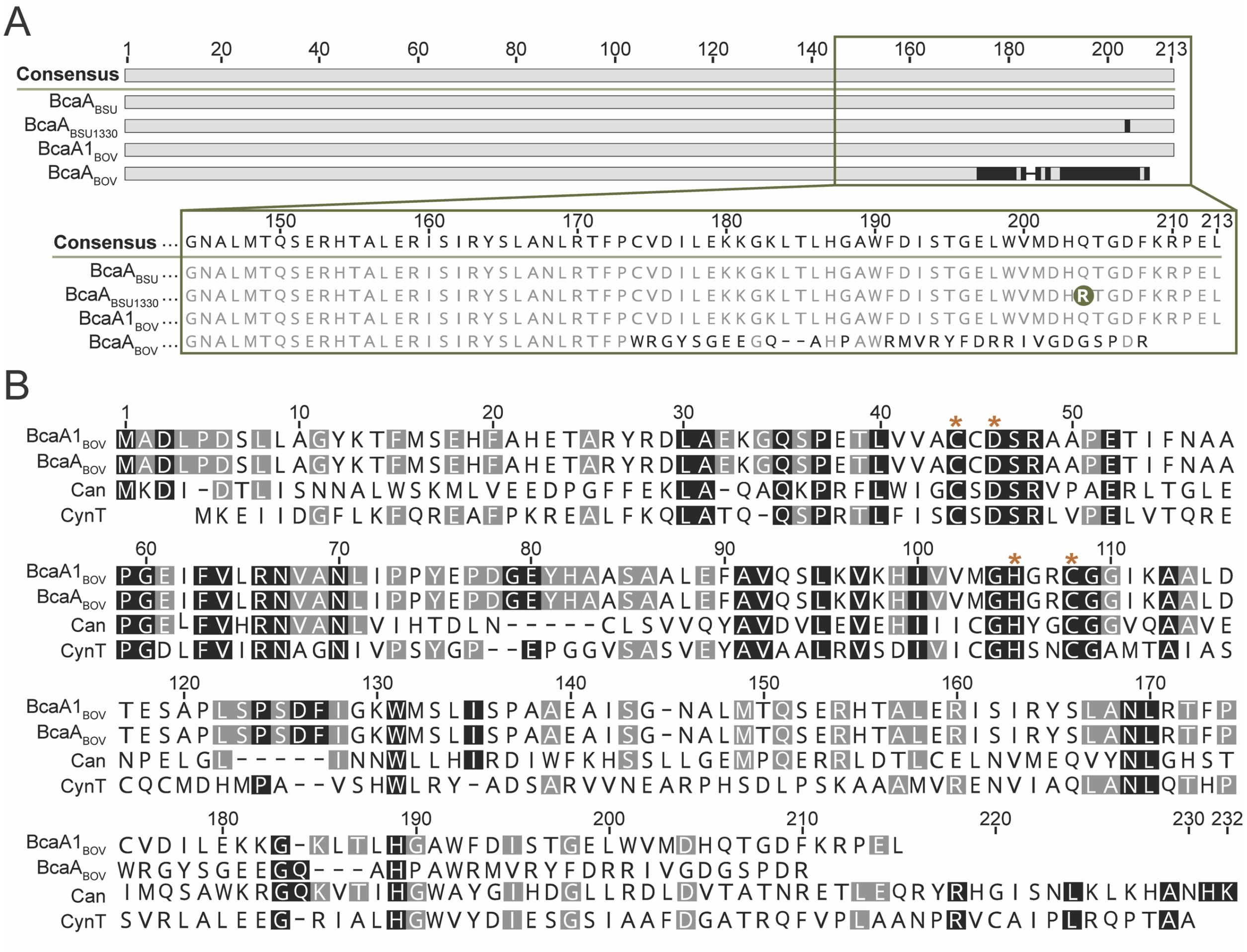


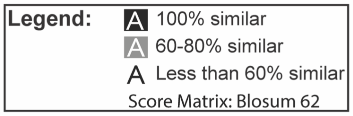


Sequence identity

| | $BcaA1_{BOV}$ | $BcaA_{BOV}$ | Can | CynT |
|---|---|---|---|---|
| $BcaA1_{BOV}$ | | 85.2% | 27.9% | 30.9% |
| $BcaA_{BOV}$ | 85.2% | | 25.7% | 28.1% |
| Can | 27.9% | 25.7% | | 27.8% |
| CynT | 30.9% | 28.1% | 27.8% | |

**Supplemental Figure 2. Comparison of *B. suis* BcaA and *E. coli* β-carbonic anhydrases to $BcaA_{BOV}$ and $BcaA1_{BOV}$**

**A) (Top)** Multiple alignment of *Brucella ovis* $BcaA_{BOV}$ against *Brucella suis* ATCC 23445 ($BcaA_{BSU}$) and *Brucella suis* 1330 ($BcaA_{BSU1330}$) homologs. The selected allele $BcaA1_{BOV}$, which enables growth of *B. ovis* without added $CO_2$, is included for comparison. Highlighted in **black** are the differences between each sequence and the genus-level consensus at **top** (see also **Fig. 3 legend**). (**Bottom**) Zoom-in on the C-terminal portion of the alignment that contains the *$bcaA_{BOV}$* frameshift present in wild-type *B. ovis*. A glutamine to arginine difference at position 204 of $BcaA_{BSU1330}$ is shaded in **green. B**) Multiple alignment of *Escherichia coli* MG1655 Can and CynT β-carbonic anhydrases with $BcaA_{BOV}$ and $BcaA1_{BOV}$. Orange asterisks show the conserved residues at the active site. **Bottom right**: table showing the amino acid identity between the proteins.

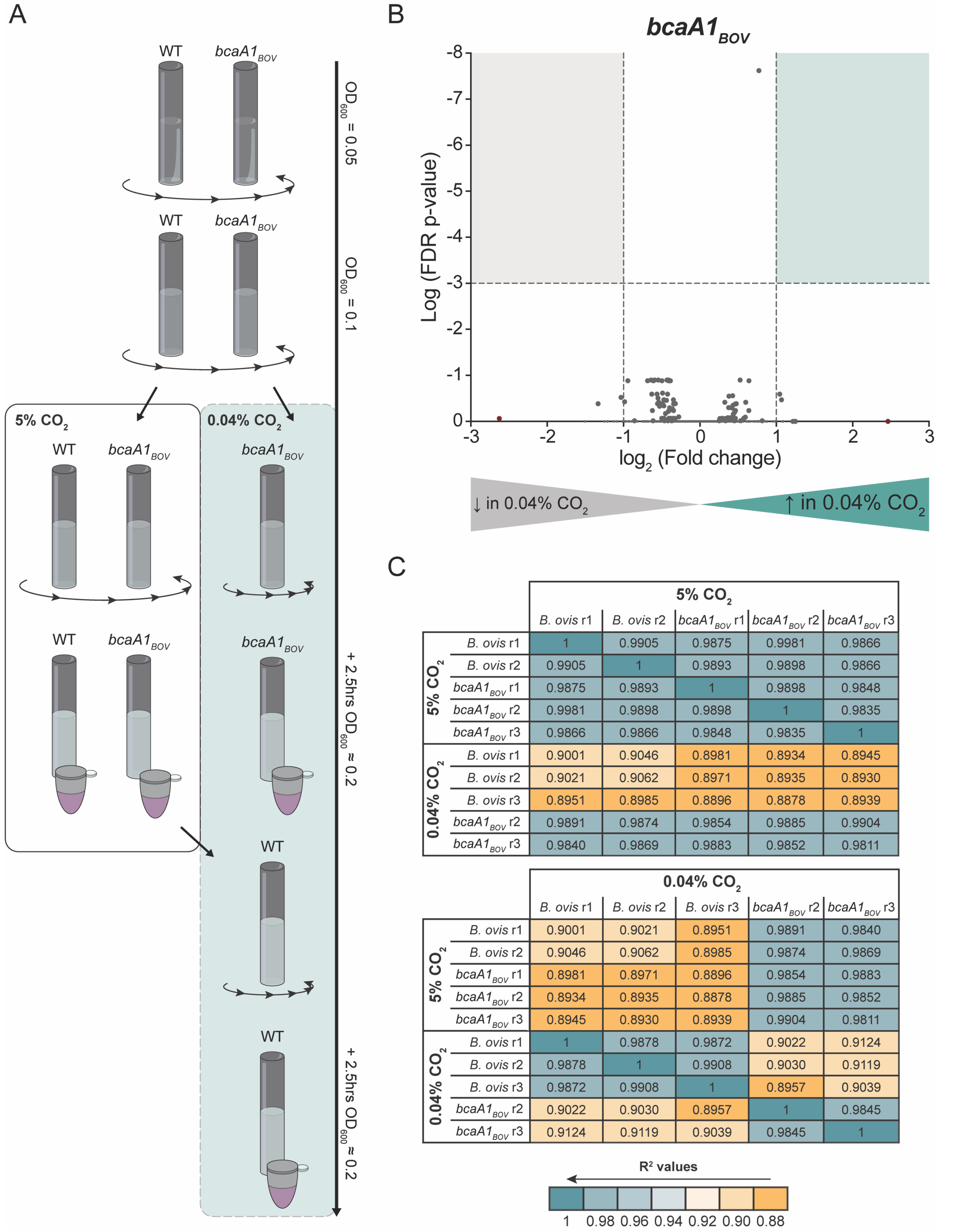


| | | 5% $CO_2$ | | | | |
|---|---|---|---|---|---|---|
| | | *B. ovis* r1 | *B. ovis* r2 | *bcaA1*$_{BOV}$ r1 | *bcaA1*$_{BOV}$ r2 | *bcaA1*$_{BOV}$ r3 |
| 5% $CO_2$ | *B. ovis* r1 | 1 | 0.9905 | 0.9875 | 0.9981 | 0.9866 |
| | *B. ovis* r2 | 0.9905 | 1 | 0.9893 | 0.9898 | 0.9866 |
| | *bcaA1*$_{BOV}$ r1 | 0.9875 | 0.9893 | 1 | 0.9898 | 0.9848 |
| | *bcaA1*$_{BOV}$ r2 | 0.9981 | 0.9898 | 0.9898 | 1 | 0.9835 |
| | *bcaA1*$_{BOV}$ r3 | 0.9866 | 0.9866 | 0.9848 | 0.9835 | 1 |
| 0.04% $CO_2$ | *B. ovis* r1 | 0.9001 | 0.9046 | 0.8981 | 0.8934 | 0.8945 |
| | *B. ovis* r2 | 0.9021 | 0.9062 | 0.8971 | 0.8935 | 0.8930 |
| | *B. ovis* r3 | 0.8951 | 0.8985 | 0.8896 | 0.8878 | 0.8939 |
| | *bcaA1*$_{BOV}$ r2 | 0.9891 | 0.9874 | 0.9854 | 0.9885 | 0.9904 |
| | *bcaA1*$_{BOV}$ r3 | 0.9840 | 0.9869 | 0.9883 | 0.9852 | 0.9811 |

| | | 0.04% $CO_2$ | | | | |
|---|---|---|---|---|---|---|
| | | *B. ovis* r1 | *B. ovis* r2 | *B. ovis* r3 | *bcaA1*$_{BOV}$ r2 | *bcaA1*$_{BOV}$ r3 |
| 5% $CO_2$ | *B. ovis* r1 | 0.9001 | 0.9021 | 0.8951 | 0.9891 | 0.9840 |
| | *B. ovis* r2 | 0.9046 | 0.9062 | 0.8985 | 0.9874 | 0.9869 |
| | *bcaA1*$_{BOV}$ r1 | 0.8981 | 0.8971 | 0.8896 | 0.9854 | 0.9883 |
| | *bcaA1*$_{BOV}$ r2 | 0.8934 | 0.8935 | 0.8878 | 0.9885 | 0.9852 |
| | *bcaA1*$_{BOV}$ r3 | 0.8945 | 0.8930 | 0.8939 | 0.9904 | 0.9811 |
| 0.04% $CO_2$ | *B. ovis* r1 | 1 | 0.9878 | 0.9872 | 0.9022 | 0.9124 |
| | *B. ovis* r2 | 0.9878 | 1 | 0.9908 | 0.9030 | 0.9119 |
| | *B. ovis* r3 | 0.9872 | 0.9908 | 1 | 0.8957 | 0.9039 |
| | *bcaA1*$_{BOV}$ r2 | 0.9022 | 0.9030 | 0.8957 | 1 | 0.9845 |
| | *bcaA1*$_{BOV}$ r3 | 0.9124 | 0.9119 | 0.9039 | 0.9845 | 1 |

**Supplemental figure 3. RNA-seq experimental set up and measured gene expression changes in *B. ovis bcaA1*$_{BOV}$ upon $CO_2$ downshift**

**A)** Schematic of sample treatment prior to RNA-seq. *B. ovis* wild-type and *bcaA1*$_{BOV}$ strains were inoculated in BB at $OD_{600}$ of 0.05 and grown with 5% $CO_2$ (6 tubes per strain, only one is shown for clarity). Once cells reached $OD_{600}$ ~0.1, 3 tubes of *B. ovis bcaA1*$_{BOV}$ cells were moved to roller in an air incubator (0.04% $CO_2$). Cells in all tubes were then grown to $OD_{600}$ ~0.2, and the three culture tubes of *Brucella ovis* ATCC 25840 (WT) in 5% $CO_2$ and all six *B. ovis bcaA1$_{BOV}$* (*bcaA1$_{BOV}$*) cultures (3 from 5% $CO_2$ and 3 from 0.04% $CO_2$) were harvested at this point for RNA prep. Three remaining WT tubes were then moved to the air incubator (0.04% $CO_2$) where they were incubated for 2.5 hours before harvesting for RNA preparation. **B)** Volcano plot for genes differentially expressed in the *B. ovis bcaA1$_{BOV}$* strain upon downshift from 5% $CO_2$ to 0.04% $CO_2$. Analysis as described for wild-type *B. ovis* ATCC 25840 in **Fig. 5B.** No genes showed significant differential expression between these two $CO_2$ conditions (FDR < 0.001 and $\log_2$ fold change > |1|). See also **Data Set 2**. **C)** $R^2$ values calculated when plotting $\log_{10}$ Counts Per Million (CPM) of *B. ovis* ATCC 25840 (WT) against *B. ovis bcaA1$_{BOV}$* in 0.04% or 5% $CO_2$. Replicates (r1-3) are indicated. Low $R^2$ values are highlighted in a **yellow** gradient, high $R^2$ values in a **teal** gradient, legend at the bottom.

**Table S1. Growth of independent spontaneous mutants derived from forward genetic selection grown in an air incubator (0.04% $CO_2$) after isolation[a]**

| Exp.[b] | Samples | $OD_{600}$ at 24hrs | $OD_{600}$ at 48hrs | WGS Samples[c] |
|---|---|---|---|---|
| Bov | WT1 | 0 | 0.001 | Parent |
| | WT2 | 0 | 0 | |
| Plate | 1 | 0.011 | 1.645 | Mutant 01 |
| liq1 | 2 | 0.026 | 1.558 | Mutant 02 |
| liq2 | 3 | 0.01 | 1.654 | Mutant 03 |
| | 4 | 0.015 | 1.664 | |
| | 5 | 0.014 | 1.654 | |
| | 6 | 0.022 | 1.655 | Mutant 14 |
| | 7 | 0.007 | 1.64 | |
| | 8 | 0.014 | 1.657 | Mutant 05 |
| | 9 | 0.005 | 1.617 | Mutant 04 |
| | 10 | 0.009 | 1.646 | |
| | 11 | 0.01 | 1.647 | |
| | 12 | 0.012 | 1.655 | Mutant 06 |
| liq3 | 13 | 0.044 | 1.633 | Mutant 07 |
| | 14 | 0.02 | 1.639 | Mutant 08 |
| | 15 | 0.018 | 1.627 | |
| | 16 | 0.027 | 1.628 | |
| | 17 | 0.012 | 1.617 | Mutant 09 |
| | 18 | 0.015 | 1.609 | |
| | 19 | 0.18 | 1.642 | Mutant 15 |
| | 20 | 0.012 | 1.629 | |
| | 21 | 0.26 | 1.606 | Mutant 10 |
| liq4 | 22 | 0.021 | 1.651 | Mutant 11 |
| | 23 | 0.024 | 1.593 | |
| | 24 | 0.026 | 1.554 | Mutant 12 |
| | 25 | 0.038 | 1.635 | Mutant 13 |
| | 26 | 0.027 | 1.635 | |
| | 27 | 0.028 | 1.635 | Mutant 16 |
| | 28 | 0.068 | 1.638 | |
| | 29 | 0.028 | 1.634 | |

[a]Cultures were inoculated at an $OD_{600}$ of $1.5 \times 10^{-5}$

[b]Spontaneous independent mutants from independent experiments (Exp.). "Plate" refers to selection conducted on solid media instead of liquid, by transferring plated cells from 5% to 0.04% $CO_2$ incubators. Liquid (liq) 1-4 are four independent experiments were cells were inoculated in broth. See **main text** and **Materials and Methods**. "Bov" is the parent strain (*Brucella ovis* ATCC 25840) where two independent tubes were inoculated for comparison with the mutants.

[c]Samples indicated were sent for whole genome sequencing (WGS).

### Table S2. Gene and pseudogene comparison across *B. ovis* and *B. abortus* ATCC 2308 strains

| Group | Gene | *B. ovis* old locus | *B. ovis* new locus | *B. abortus* locus | Gene length[a] | Polymorphisms between *B. ovis* and *B. abortus*[b] | | | | Polymorphisms within *B. ovis* genomes | |
|---|---|---|---|---|---|---|---|---|---|---|---|
| | | | | | | SNPs | | Single nt Indels | Indels >1nt | # site(s)[d] | # genomes[e] |
| | | | | | | Total | Syn[c] | | | | |
| Pseudogenes | *ureF2* | BOV_1316 | BOV_RS06515 | BAB_RS22515 | 732 | 3 | / | 2 | 0 | 0 | 0 |
| | *ureT* | BOV_1319 | BOV_RS06530 | BAB_RS22530 | 1050 | 2 | / | 0 | 1 (56nt) | 1 | 15 |
| | *ureG1* | BOV_0287 | BOV_RS01465 | BAB_RS17375 | 627 | 1 | / | 0 | 0 | 0 | 0 |
| | *ureE2* | BOV_1315 | BOV_RS06510 | BAB_RS22510 | 606 | 2 | / | 2 | 0 | 0 | 0 |
| | *ureC1* | BOV_0284 | BOV_RS01450 | BAB_RS17360 | 1713 | 5 | / | 0 | 1 (30nt) | 0 | 0 |
| | *pckA* | BOV_2009 | BOV_RS09880 | BAB_RS25895 | 1476 | 8 | / | 1 | 0 | 1 | 1 |
| | *eryA* | BOV_A0811 | BOV_RS14445 | BAB_RS28140 | 1554 | 6 | / | 0 | 0 | 0 | 0 |
| | *eryD* | BOV_A0814 | BOV_RS14460 | BAB_RS28125 | 951 | 4 | / | 0 | 1 (7nt) | 0 | 0 |
| | *eryF* | BOV_A0805 | BOV_RS14425 | BAB_RS28160 | 948 | 4 | / | 0 | 1 (9nt) | 0 | 0 |
| | *eryG* | BOV_A0806 | BOV_RS14430 | BAB_RS28155 | 1041 | 1 | / | 0 | 1 (2nt) | 0 | 0 |
| | *gluP* | BOV_A0172 | BOV_RS11215 | BAB_RS27245 | 1239 | 3 | / | 1 | 0 | 0 | 0 |
| | *ccoO* | BOV_0378 | BOV_RS01915 | BAB_RS17800 | 732 | 1 | / | 1 | 0 | 1 | 11 |
| | *coxB* | / | BOV_RS02370 | BAB_RS18265 | 889 | 2 | / | 1 | 0 | 0 | 0 |
| | *coxG* | BOV_0478 | BOV_RS16275 | BAB_RS18290 | 879 | 2 | / | 1 | 2 (56nt, 6nt) | 0 | 0 |
| | *ctaE* | / | BOV_RS11490 | BAB_RS30835 | 570 | 2 | / | 0 | 2 (12nt, 27nt) | 0 | 0 |
| | *ctaG* | BOV_0477 | BOV_RS02390 | BAB_RS18280 | 606 | 1 | / | 0 | 1 (5nt) | 0 | 0 |
| | *norB* | / | BOV_RS11505 | BAB_RS30820 | 1350 | 6 | / | 0 | 1 (80nt) | 1 | 1 |
| | *copA/fixI* | / | BOV_RS01885 | BAB_RS17775 | 2259 | 5 | / | 0 | 1 (153nt) | 0 | 0 |
| | *bcaA* | / | BOV_RS08635 | BAB_RS24650 | 645 | 0 | / | 1 | 1 (2nt) | 0 | 0 |
| Urease genes (not pseudogenes) | *ureD1* | BOV_0281 | BOV_RS01430 | BAB_RS17340 | 843 | 2 | 1 | 0 | 0 | 0 | 0 |
| | *ureA1* | BOV_0202 | BOV_RS01435 | BAB_RS17345 | 303 | 1 | 0 | 0 | 0 | 0 | 0 |
| | *ureB1* | BOV_0283 | BOV_RS01445 | BAB_RS17355 | 306 | 0 | 0 | 0 | 0 | 0 | 0 |
| | *ureE1* | BOV_0285 | BOV_RS01455 | BAB_RS17365 | 516 | 5 | 3 | 0 | 0 | 0 | 0 |
| | *ureF1* | BOV_0286 | BOV_RS01460 | BAB_RS17370 | 688 | 2 | 0 | 0 | 0 | 1 | 1 |
| | *ureA2* | BOV_1312 | BOV_RS06495 | BAB_RS22495 | 303 | 1 | 0 | 0 | 0 | 0 | 0 |
| | *ureB2* | BOV_1313 | BOV_RS06500 | BAB_RS22500 | 480 | 2 | 0 | 0 | 0 | 0 | 0 |
| | *ureC2* | BOV_1314 | BOV_RS06505 | BAB_RS22505 | 1722 | 5 | 1 | 0 | 0 | 0 | 0 |
| | *ureG2* | BOV_1381 | BAB_RS22520 | BAB_RS22520 | 639 | 1 | 1 | 0 | 0 | 0 | 0 |
| | *ureD2* | BOV_1318 | BOV_RS06525 | BAB_RS22525 | 915 | 4 | 1 | 0 | 1 (6nt) | 0 | 0 |
| Type IV Secretion System | *virB11* | BOV_A0054 | BOV_RS10610 | BAB_RS26635 | 1086 | 2 | 2 | 0 | 0 | 0 | 0 |
| | *virB10* | BOV_A0055 | BOV_RS10615 | BAB_RS26640 | 1167 | 3 | 1 | 0 | 1 (24nt) | 1 | 1 |
| | *virB9* | BOV_A0056 | BOV_RS10620 | BAB_RS26645 | 870 | 0 | 0 | 0 | 0 | 1 | 1 |
| | *virB8* | BOV_A0057 | BOV_RS10625 | BAB_RS26650 | 720 | 2 | 1 | 0 | 0 | 0 | 0 |
| | *virB7* | / | BOV_RS10630 | BAB_RS26655 | 174 | 0 | 0 | 0 | 0 | 0 | 0 |
| | *virB6* | BOV_A0058 | BOV_RS10635 | BAB_RS26660 | 1044 | 1 | 1 | 0 | 0 | 1 | 11 |
| | *virB5* | BOV_A0059 | BOV_RS10640 | BAB_RS26665 | 717 | 4 | 2 | 0 | 0 | 0 | 0 |
| | *virB4* | BOV_A0060 | BOV_RS10645 | BAB_RS26670 | 2496 | 4 | 1 | 0 | 0 | 2 | 2[f] |
| | *virB3* | BOV_A0061 | BOV_RS10650 | BAB_RS26675 | 351 | 1 | 1 | 1 | 0 | 0 | 0 |
| | *virB2* | BOV_A0062 | BOV_RS10655 | BAB_RS26680 | 318 | 0 | 0 | 0 | 0 | 0 | 0 |
| | *virB1* | BOV_A0063 | BOV_RS10660 | BAB_RS26685 | 717 | 2 | 0 | 0 | 0 | 0 | 0 |

[a] In *B. abortus* ATCC 2308
[b] Number of polymorphisms between all 17 sequenced *B. ovis* clinical isolates (see **Data Sheet 1**) and *B. abortus* ATCC 2308 strain (GenBank accessions NC_007618 and NC_007624).
[c] Syn = synonymous mutations. In the case of pseudogenes, synonymous mutations were not noted.
[d] Number of polymorphic sites among sequences from *B. ovis* isolates
[e] Number of *B. ovis* sequences that harbor the polymorphism
[f] Two genomes have distinct polymorphisms
